# Skeletal muscle methylome and transcriptome integration reveals profound sex differences related to muscle function and substrate metabolism

**DOI:** 10.1101/2021.03.16.435733

**Authors:** Shanie Landen, Macsue Jacques, Danielle Hiam, Javier Alvarez-Romero, Nicholas R Harvey, Larisa M. Haupt, Lyn R Griffiths, Kevin J Ashton, Séverine Lamon, Sarah Voisin, Nir Eynon

**Author notes:** Sarah Voisin and Nir Eynon are co-senior authors. **Corresponding author:** Professor Nir Eynon, Institute for Health and Sport (iHeS), Victoria University, PO Box 14428, Melbourne, VIC 8001, Australia.

## Abstract

Nearly all human complex traits and diseases exhibit some degree of sex differences, with epigenetics being one of the main contributing factors. Various tissues display sex differences in DNA methylation, however this has not yet been explored in skeletal muscle, despite skeletal muscle being among the tissues with the most transcriptomic sex differences. For the first time, we investigated the effect of sex on autosomal DNA methylation in human skeletal muscle across three independent cohorts (Gene SMART, FUSION, and GSE38291) using a meta-analysis approach, totalling 369 human muscle samples (222 males, 147 females), and integrated this with known sex-biased transcriptomics. We found 10,240 differentially methylated regions (DMRs) at FDR < 0.005, 94% of which were hypomethylated in males, and gene set enrichment analysis revealed that differentially methylated genes were involved in muscle contraction and substrate metabolism. We then investigated biological factors underlying DNA methylation sex differences and found that circulating hormones were not associated with differential methylation at sex-biased DNA methylation loci, however these sex-specific loci were enriched for binding sites of hormone-related transcription factors (with top TFs including androgen (*AR*), estrogen (*ESR1*), and glucocorticoid (*NR3C1*) receptors). Fibre type proportions were associated with differential methylation across the genome, as well as across 16 % of sex-biased DNA methylation loci (FDR < 0.005). Integration of DNA methylomic results with transcriptomic data from the GTEx database and the FUSION cohort revealed 326 autosomal genes that display sex differences at both the epigenome and transcriptome levels. Importantly, transcriptional sex-biased genes were overrepresented among epigenetic sex-biased genes (p-value = 4.6e-13), suggesting differential DNA methylation and gene expression between male and female muscle are functionally linked. Finally, we validated expression of three genes with large effect sizes (*FOXO3A, ALDH1A1*, and *GGT7*) in the Gene SMART cohort with qPCR. *GGT7*, involved in antioxidant metabolism, displays male-biased expression as well as lower methylation in males across the three cohorts. In conclusion, we uncovered 8,420 genes that exhibit DNA methylation differences between males and females in human skeletal muscle that may modulate mechanisms controlling muscle metabolism and health.

**Significance:** The importance of uncovering biological sex differences and their translation to physiology has become increasingly evident. Using a large-scale meta-analysis of three cohorts, we perform the first comparison of genome-wide skeletal muscle DNA methylation between males and females, and identify thousands of genes that display sex-differential methylation. We then explore intrinsic biological factors that may be underlying the DNA methylation sex differences, such as fibre type proportions and sex hormones. Leveraging the GTEx database, we identify hundreds of genes with both sex-differential expression and DNA methylation in skeletal muscle. We further confirm the sex-biased genes with gene expression data from two cohorts included in the methylation meta-analysis. Our study integrates genomewide sex-biased DNA methylation and expression in skeletal muscle, shedding light on distinct sex differences in skeletal muscle.

## Introduction

Sex differences are evident in nearly all complex traits. Various diseases, including but not limited to cancer, muscular dystrophy, and COVID-19 [3, 4], display sex differences in prevalence, onset, progression, or severity. To improve treatment for such diseases, it is crucial to uncover the molecular basis for the sex differences and their consequences on organ function [5]. Sexually differentiated traits and phenotypes stem from a combination of factors, including genetics (gene variants-by-sex interactions [6], XY chromosome complements [7–10], genomic imprinting [11]), the hormonal milieu [12, 13], and gene regulation [14], with the latter likely contributing the most [3].

Recently, a large-scale study from the Genotype-Tissue Expression (GTEx) consortium unravelled mRNA expression differences between the sexes that are not driven by sex chromosomes, across all tissues. Skeletal muscle was particularly divergent between the sexes, as gene expression profiles in this tissue could predict sex with high specificity and sensitivity [7]. These transcriptomic differences underpin the numerous physiological differences in skeletal muscle between males and females, such as differences in substrate metabolism [15–17]. For example, females oxidize more lipids and less carbohydrates and amino acids during endurance exercise, and albeit depending on training status, tend to have a higher proportion of type I (slow-twitch) muscle fibres [18], all of which inherently contribute to enhanced fatigue-resistance in female skeletal muscle [19]. As such, females exhibit higher mRNA and protein levels of lipid oxidation-related genes than males [16]. Interestingly, the top gene set corresponding to sex-biased genes in the GTEx study corresponded to targets of the epigenetic writer polycomb repressive complex 2 (PRC2) and its associated epigenetic mark (H3K27me3) [7]. This suggests that the sex-specific deposition of epigenetic marks may be one of the sources of sex differences in gene expression. Epigenetics is a regulatory system that influences gene expression and is modulated by the genetic sequence and environmental stimuli. DNA methylation is currently the best-characterized epigenetic modification, and has been shown to differ between males and females in various tissues, such as pancreatic islets [20], blood [21, 22], and more recently cultured myoblasts and myotubes [23]. While there is ample evidence for transcriptomic sex differences in skeletal muscle [7, 14, 15, 24–26], it is unclear whether sex differences exist in the DNA methylome of skeletal muscle tissue, and what factors contribute to these differences. Epigenome-wide association studies (EWAS) are ideal for investigating the impact of sex on genome-wide DNA methylation profiles.

Differences in skeletal muscle fibre-type proportions and circulating levels of sex hormones may contribute to epigenomic and transcriptomic differences between the sexes. Males and females are exposed to differing levels of sex hormones across the lifespan [35], which are primarily ascribed to the reproductive function, but their importance to non-reproductive functions, such as skeletal muscle [27], is becoming more apparent [28, 29]. Sex steroid hormones engage through their specific ligand-receptors [7, 30, 31] and influence transcription and phenotype in a tissue- and sex-specific manner [12, 32–35]. Not only can these receptors be differentially expressed between sexes [36], they also show sex-biased gene targeting patterns due to intrinsic differences in sex hormone levels [31, 37]. Human muscle consists of three distinct muscle fibre types; type I (slow-twitch oxidative), type IIa (fast-twitch oxidative), and type IIx (fast-twitch) fibres. Males tend to have a higher proportion of fast-twitch type IIa muscle fibres in various muscle groups compared to females [38, 39]. The different fibre types exhibit different methylation patterns [40], as well as different contractile and metabolic properties [41]. Thus, differences in fibre type proportions may underlie sex differences in skeletal muscle DNA methylation.

We performed a large-scale EWAS meta-analysis to explore sex differences in the DNA methylome of human skeletal muscle tissue, using three datasets from our own laboratory and open-access databases (n = 369 individuals; 217 males, 152 females). We established a list of sites (CpG) and regions showing DNA methylation differences between males and females, and explored their genomic context. We then investigated whether transcription factor binding, muscle fibre type distribution, and circulating sex hormone levels explained the observed sex differences in DNA methylation. Next, we integrated the sex-biased DNA methylation with known sex-biased mRNA expression from the GTEx consortium, and inferred the potential downstream effects on skeletal muscle function. Lastly, we validated our findings with transcriptomic data from one cohort used in the meta-analysis and targeted qPCR from another cohort.

## Results

### Males show profound genome-wide autosomal hypomethylation compared with females in human skeletal muscle

The DNA methylation meta-analysis was conducted on 369 individuals from three datasets (217 males, 152 females). We focused exclusively on the 22 autosomes as sex chromosomes represent a particular case whereby epigenetically-driven X-chromosome inactivation takes place exclusively in females. All of the Gene SMART participants were apparently healthy, while the FUSION participants were either healthy or diagnosed with type 2 diabetes mellitus (T2D), and the GSE38291 consisted of monozygotic twins discordant for T2D (**Table 1**).

**Table 1.** Characteristics of participants in each data set included in the DNA methylation meta-analysis. Statistics shown for differences between males and females.

| FUSION | Females, N = 115<sup>1</sup> | Males, N = 159<sup>1</sup> | p-value<sup>2</sup> | Gene SMART | Females, N = 20<sup>1</sup> | Males, N = 45<sup>1</sup> | p-value<sup>2</sup> | GSE38291 | Females, N = 12<sup>1</sup> | Males, N = 10<sup>1</sup> | p-value<sup>2</sup> |
| --- | --- | --- | --- | --- | --- | --- | --- | --- | --- | --- | --- |
| Age (years) | 61 (8) | 59 (8) | 0.2 | Age (years) | 35 (7) | 32 (8) | 0.10 | Age (years) | 66 (9) | 70 (4) | 0.15 |
| Health |  |  | 0.026 | Health |  |  |  | Health |  |  |  |
| Healthy | 93 (81%) | 109 (69%) |  | Healthy | 20 (100%) | 45 (100%) |  | Healthy | 6 (50%) | 5 (50%) |  |
| T2D | 22 (19%) | 50 (31%) |  |  |  |  |  | T2D | 6 (50%) | 5 (50%) |  |
| <sup>1</sup> Mean (SD); n (%) |  |  |  | <sup>1</sup> Mean (SD); n (%) |  |  |  | <sup>1</sup> Mean (SD); n (%) |  |  |  |
| <sup>2</sup> Welch Two Sample t-test; Fisher's exact test |  |  |  | <sup>2</sup> Welch Two Sample t-test |  |  |  | <sup>2</sup> Welch Two Sample t-test |  |  |  |

We found 56,813 differentially methylated positions (DMPs, single CpG sites) between males and females, spread across the 22 autosomes, at a meta-analysis false discovery rate (FDR) < 0.005 (**Figure 1**, Supplementary Table 2). Ninety-four percent of DMPs were hypomethylated in males compared with females (**Figure 1A**). On average, DNA methylation levels differed by +2.8% (hyper DMPs) and −3.5% (hypo DMPs) between males and females, with the largest effect sizes reaching +15.2% and −35.7%. While participants did not cluster according to sex when looking at the whole methylome, they did cluster according to sex when only focusing on the 56,813 DMPs (**Figure 1B**), suggesting that sex explained a substantial amount of variance at the DMPs.

**Figure 1.**
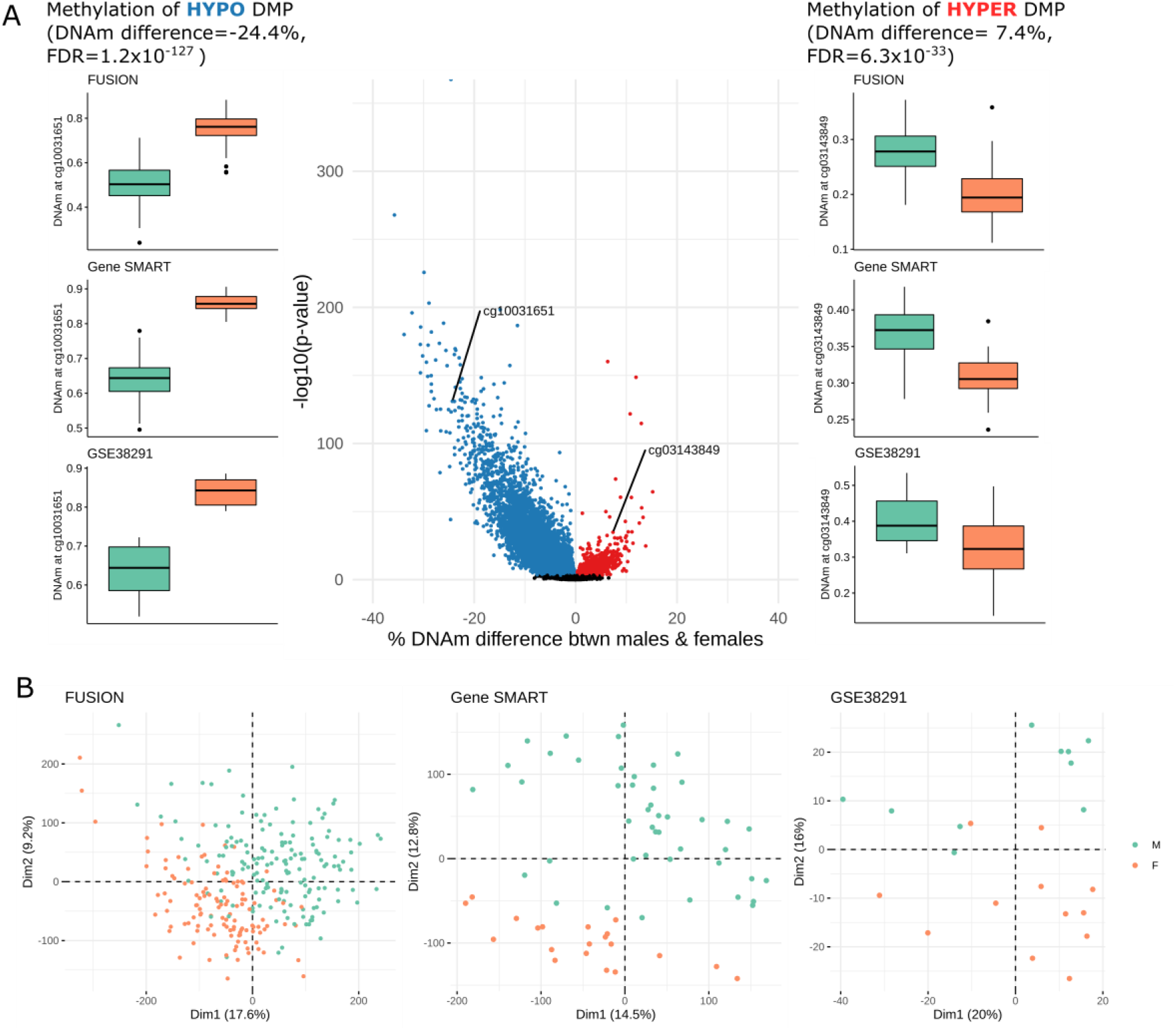
Differentially methylated positions (DMPs) with sex in skeletal muscle. (**A**)Volcano plot of DNA methylation differences between males and females. Each point represents a tested CpG (633,645 in total) and those that appear in color are DMPs at a meta-analysis false discovery rate < 0.005; red DMPs are hypermethylated in males compared with females; blue DMPs are hypomethylated in males compared with females. The x-axis represents the amount of DNA methylation difference between the sexes and the y-axis represents statistical significance (higher = more significant). Two DMPs that were present in all three studies and showed the largest effect size are labeled with the respective CpG and boxplots of β-values from each study appear to the right (hyper DMP) and left (hypo DMP). (**B**) Principal component analysis plots of the methylation values at the DMPs; each point on the graph represents an individual; males denoted in green, females denoted in orange.

Each data set had a unique study design that required adjustment for factors known to affect DNA methylation, such as age [42] and type 2 diabetes (T2D) [43]. We adjusted each dataset for these factors, but noted that sex was associated with T2D in the FUSION dataset, meaning that male participants from the FUSION cohort more commonly had T2D than females. Therefore, it is possible that the sex-related signal captured in this dataset was partially confounded by T2D. We repeated the meta-analysis excluding T2D participants from the FUSION cohort, but results remained unchanged (Supplementary Figure 7).

### Fibre type proportions, but not circulating hormones, were associated with differential methylation at loci with sex-specific DNA methylation

Males typically show a greater proportion of type II muscle fibres compared with females [18], and type II fibres exhibit hypomethylation compared to type I fibres [40]. Therefore, we hypothesized that the observed DNA methylation sex differences, specifically the hypomethylation in males, may be a result of differing fibre type distributions between males and females. We first estimated type I fibre proportions in the Gene SMART cohort via immunohistochemistry (Supplementary Figure 2D) and the FUSION cohort via RNA-seq (data available for each dataset in Supplementary Table 13). In both the Gene SMART and FUSION cohorts, females had higher proportions of type I fibres than males (Supplementary Figure 2B/C). We could not directly add fibre type proportions to the linear model as a covariate, since fibre type proportions are not a *confounder* (i.e. a factor that influences both sex and DNA methylation independently), but may be a direct *downstream effect* of sex, in turn affecting DNA methylation. Adding fibre type proportions in the model would therefore distort the association between sex and DNA methylation. To overcome this issue, we stratified the cohorts by sex, added fibre type proportions to the model as a covariate and identified DNA methylation patterns associated with fibre type proportions. We then meta-analysed the results to find CpGs robustly associated with fibre type proportions across both cohorts and all sexes (see “Methods”). We identified 16,275 CpGs associated with fibre type proportions (Supplementary Figure 2). When restricting the analysis to the loci exhibiting sex-biased DNA methylation, 8,805 (15.5%) of those were associated with fibre type proportions (FDR < 0.005). Effect sizes ranged from −0.28% to +0.30% DNA methylation difference per % increase in type I fibre content (**Figure 2A**).

**Figure 2.**
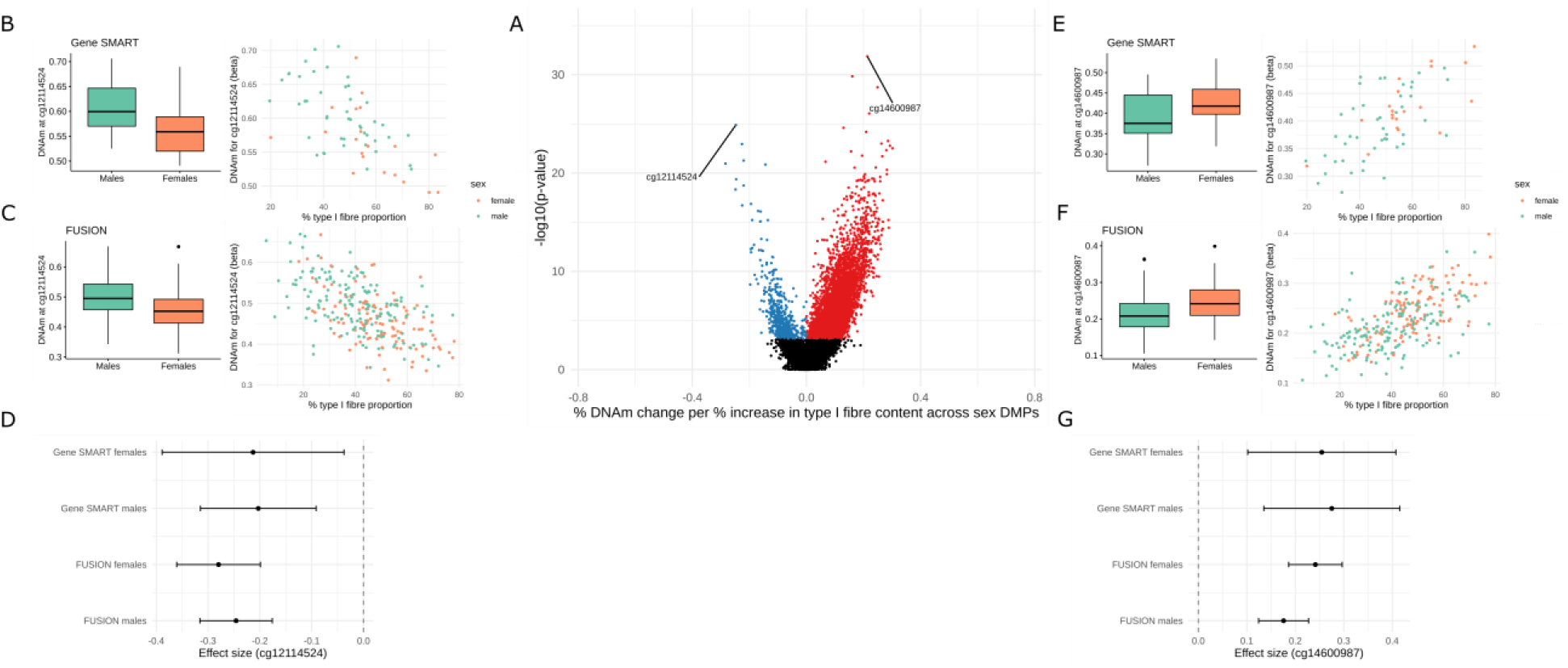
Fibre type-related DNA methylation loci across sex-biased DNA methylation loci. (**A**) Meta-analysis effect size (*x-axis*) and meta-analysis significance (*y-axis*) for the 56,813 tested sex-biased CpGs. Hypomethylated (blue) and hypermethylated (red) point represent differentially methylated positions (DMPs) at false discovery rate (FDR) < 0.005. One hyper- and one hypo-DMP which showed the largest effect sizes are labeled with the respective CpG; with boxplots of β-values per sex and scatter plots of β-values relative to type I fibre proportion from the Gene SMART (**B,C**) and FUSION (**E,F**) cohorts. Females are represented in orange and males in green. (**D,G**) Forest plots for the given CpG, showing effect size and confidence intervals for each sex in each study.

We then aimed to determine whether circulating sex hormone levels underlie the observed DNA methylation sex differences. We analysed oestrogen (as oestradiol, E2), testosterone (T), free testosterone (Free T), and sex-hormone binding globulin (SHBG) levels using mass-spectrometry (Free T derived from calculation) of blood serum in males and females in the Gene SMART cohort. Males and females significantly differed in all four hormone levels (Supplementary Table 12). To avoid collinearity with sex, we separated males and females for the association between sex-differential DNA methylation and circulating hormones. We assessed whether each of the four hormone levels was associated with DNA methylation across all of the CpGs and across the sex-DMPs in each sex by adjusting the linear model for a given hormone. In both males and females, circulating free testosterone, testosterone, oestrogen, and SHBG levels were not highly associated with DNA methylation (less than five DMPs; FDR < 0.005) of neither all of the CpGs tested nor the sex-DMPs previously identified (Supplementary Figure 3). To limit the potentially confounding effect of fluctuating ovarian hormone levels on DNA methylation, female muscle biopsies were collected in the early follicular phase of the menstrual cycle (days 1-7) and blood serum were tested for follicle stimulating hormone (FSH), luteinizing hormone (LH), and progesterone (as well as E2 as previously mentioned). None of the four hormones (when adjusting for each hormone separately or when adjusting for the first two principal components; see ‘methods’), were associated with differential methylation of all the CpGs or the sex-DMPs. This suggests that variations in ovarian hormone levels in the early follicular phase did not confound our results.

### Males show profound genome-wide autosomal hypomethylation compared with females in human skeletal muscle

Since the association between DNA methylation and gene expression depends on the genomic context, we inspected the genomic distribution of the DMPs to gain insights into their potential function [44]. We compared the distribution of hyper-, hypo-, and non-DMPs among the various chromatin states in human skeletal muscle using the Roadmap Epigenomics project [45]. DMPs were not randomly distributed in the chromatin states (χ^2^ p-value < 2.2 x 10^−16^, **Figure 2A**); specifically, hypo DMPs were enriched in enhancers and depleted in transcription start sites (Supplementary Figure 1A), while hyper DMPs were not enriched in any chromatin states given their scarcity. It should be noted that the Roadmap Epigenomics project characterizes both male and female skeletal muscle chromatin states regions, and there are 536 regions across 369 unique genes where male and female chromatin states differ (across many tissues including skeletal muscle) [46]. Therefore, we performed the chromatin state enrichment analysis on both the male and female chromatin state annotation in skeletal muscle, which yielded equivalent findings. We noted that DMPs were overrepresented in loci whose chromatin states differ between males and females (38.7 % of DMPs vs. 32.4% of non-DMPs are in chromatin states that differ between males and females), which means that the odds of a DMP being located in a sex-differing chromatin state increased by a factor of 1.3 compared with a non-DMP (OR = 0.76, 95% confidence interval = 0.75-0.77, Fisher test p-value < 2.2e-16) (**Figure 2B**). DMPs were also enriched in CpG island shores and depleted in CpG islands (χ^2^ p-value < 2.2e-16) (**Figure 2C**, Supplementary figure 1B).

Given the role of transcription factors (TFs) in regulating chromatin accessibility and thus effecting downstream gene expression [47], as well as the recent studies identifying sex differences in TF targeting patterns [7, 14]; we next tested whether DMPs were enriched for the experimentally validated binding sites (TFBSs) of 268 TFs from 518 different cell and tissue types [2, 48]. The DMPs were overrepresented in the binding sites of 41 TFs (p-value < 0.005, **Figure 2D, Supplementary table 14**), including hormone-related TFs such as androgen (*AR*), estrogen (*ESR1*), and glucocorticoid (*NR3C1*) receptors.

**Figure 2.**
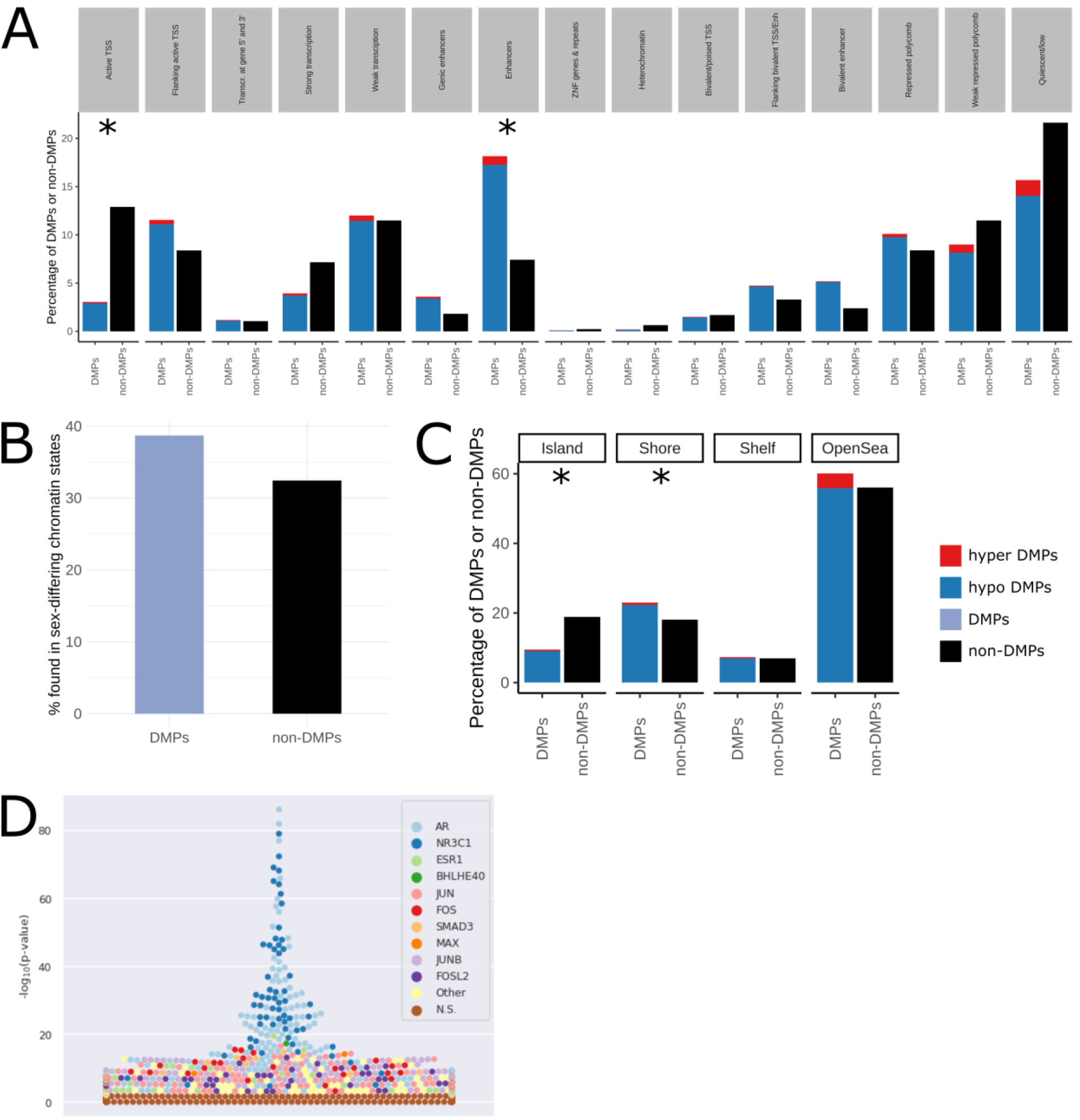
Genomic context of sex-differentially methylated positions. (**A**) Distribution of hyper/hypo DMPs and non-DMPs with respect to chromatin states (male skeletal muscle annotation). Blue is hypomethylated in males and red is hypermethylated in males. Red and blue add up to all of the sex-DMPs. Black denotes the rest of the CpG sites from the analysis which are not DMPs. Asterisks represent a greater contribution to the significant relationship between DMP status and chromatin state (Supplementary figure 1A). (**B**) Distribution of sex-DMPs and non-DMPs at loci whose chromatin states differ between male and female skeletal muscle. Purple denotes all DMPs (hypo and hyper combined) and black denotes non-DMPs. (**C**) Distribution of sex-DMPs and non-DMPs in relation to CpG islands. Asterisks represent a greater contribution to the significant relationship between DMP status and CpG island location (Supplementary figure 1B). (**D**) Bee swarm plot showing enrichment of transcription factor binding sites (TFBSs)(−(log_10_(*p* value) using Fisher’s exact tests) on the *y*-axis for differentially methylated positions (DMPs) according to UniBind [2]. The names of the top 10 enriched TFs are denoted by the colour key; brown denotes non-significant TFs. The various data sets for the same TFs are graphed with the corresponding colour.

### Genes with sex-biased methylation exhibit sex-biased DNA expression in human skeletal muscle

Differentially methylated genes (DMGs) were determined by identifying differentially methylated regions (DMRs), as DMRs remove spatial redundancy (CpG sites ~500 bp apart are typically highly correlated [49]), and may provide more robust and functionally important information than DMPs [50, 51]. We identified 10,240 DMRs (Stouffer, harmonic mean of the individual component FDRs (HMFDR), and Fisher p-value < 0.005). These DMRs were annotated to 8,420 unique autosomal genes (including non-coding genes) (Supplementary Table 3).

To gain insights into the potential downstream effects of sex-biased DNA methylation on gene expression, we integrated results from the EWAS meta-analysis of sex with genes whose mRNA expression levels are known to differ between males and females. We used version 8 of the Genotype-Tissue Expression (GTEx) database which contains 803 RNA-sequencing profiles in human skeletal muscle (n = 543 males and n = 260 females). There were 2,689 sex-differentially expressed genes (DEGs) on the autosomes in skeletal muscle (accessed from GTEx portal on 08/26/2020). Of the 2,689 DEGs, 973 (~36%) were in common with DMGs from our cohorts (**Figure 3**, Supplementary table 2), including the gene Gamma-Glutamyltransferase 7 (*GGT7*) (**Figure 5**). We confirmed an enrichment of DMRs across sex-biased genes (hypergeometric test p-value = 4.6e-13), suggesting that the overlap between sex-differentially methylated genes and sex-differentially expressed genes is larger than what would be expected by chance alone. To gain insight on the relationship between DNA methylation and gene expression of sex-biased genes, we assessed the direction of correlation between DMRs that are annotated to either promoter (TssA and TssAFlnk) or enhancer (Enh and EnhG) regions and their given gene expression (**Figure 3C-D**). Sixty-two and 59 % of DMRs in promoter and enhancer regions, respectively, were inversely correlated with gene expression (from GTEx transcriptome data, similar results were yielded with the FUSION transcriptome data). The inverse correlation between DNA methylation at both promoter and enhancer regions with gene expression was more than would be expected to occur by random chance (10,000 random permutations; p-value <0.0001 and p-value = 0.0009, respectively; Supplementary Figure 5).

**Figure 3.**
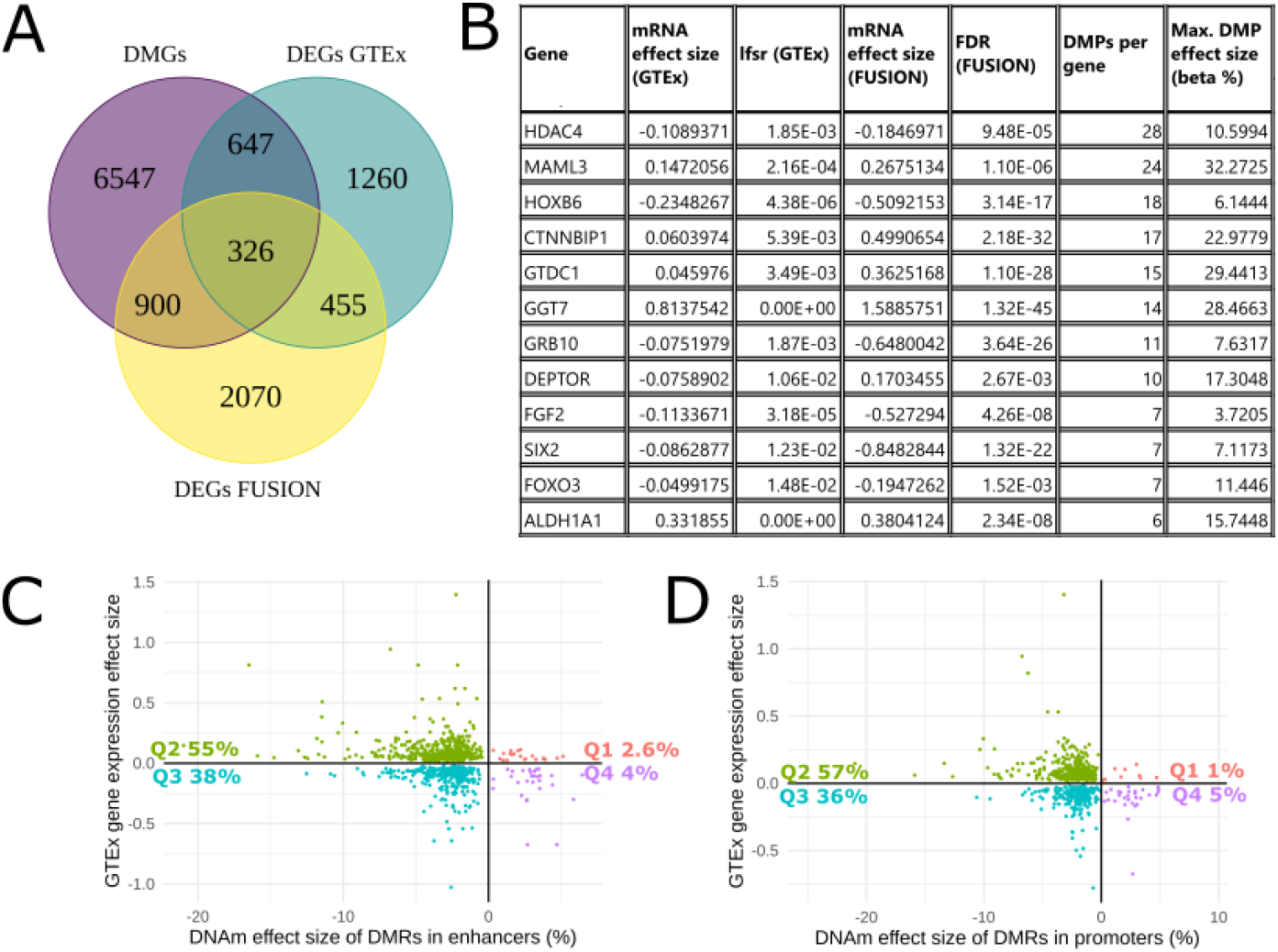
Integration of differentially methylated genes and differentially expressed genes. (**A**) Venn diagram of the overlap between differentially methylated genes (DMGs; derived from DMRs), differentially expressed genes derived from GTEx (DEGs GTEx), and differentially expressed genes derived from FUSION (DEGs FUSION) between males and females. (**B**) Subset of 12 genes with consistently large effect sizes or of biological relevance to skeletal muscle. (**C**) Correlation between the effect sizes of DMRs in enhancer regions and the effect sizes of gene expression of the relative annotated gene (for GTEx sex-biased genes). Quadrant percentages indicate the percentage DMRs/DEGs that fall into each quadrant. (**D**) Correlation between the effect sizes of DMRs in promoter regions and the effect sizes of gene expression of the relative annotated gene (for GTEx sex-biased genes). Quadrant percentages indicate the percantage DMRs/DEGs that fall into each quadrant.

### Gene set enrichment analysis of differentially methylated regions

We next performed Gene set enrichment analysis (GSEA) on the DMGs, as GSEA using epigenomic features may reveal distinct enriched pathways that may not display gene expression differences [14, 46]. We performed GSEA on both the DMRs and DMPs (**Figure 4**). GSEA on the DMRs revealed enrichment of several Gene Ontology (GO) terms, one Reactome pathway (“muscle contraction”), but no Kyoto Encyclopedia of Genes and Genomes (KEGG) pathways (Supplementary Table 10) (FDR < 0.005). However, GSEA on the DMPs revealed enrichment across all three databases (Supplementary Tables 5, 7, and 9). Most of the enriched GO terms are biological process (BP) terms, many of which relate to anatomical structure development as well as many muscle-related processes. Furthermore, DMPs were enriched for GO terms related to substrate metabolism such as lipid, protein, and carbohydrate derivative metabolic processes. Nine-hundred and twenty-five genes of the 1,407 genes involved in KEGG metabolic pathways were differentially methylated, representing many aspects of substrate metabolism (Supplementary Figure 4), although the pathway was only significant when analysing the DMPs.

**Figure 4.**
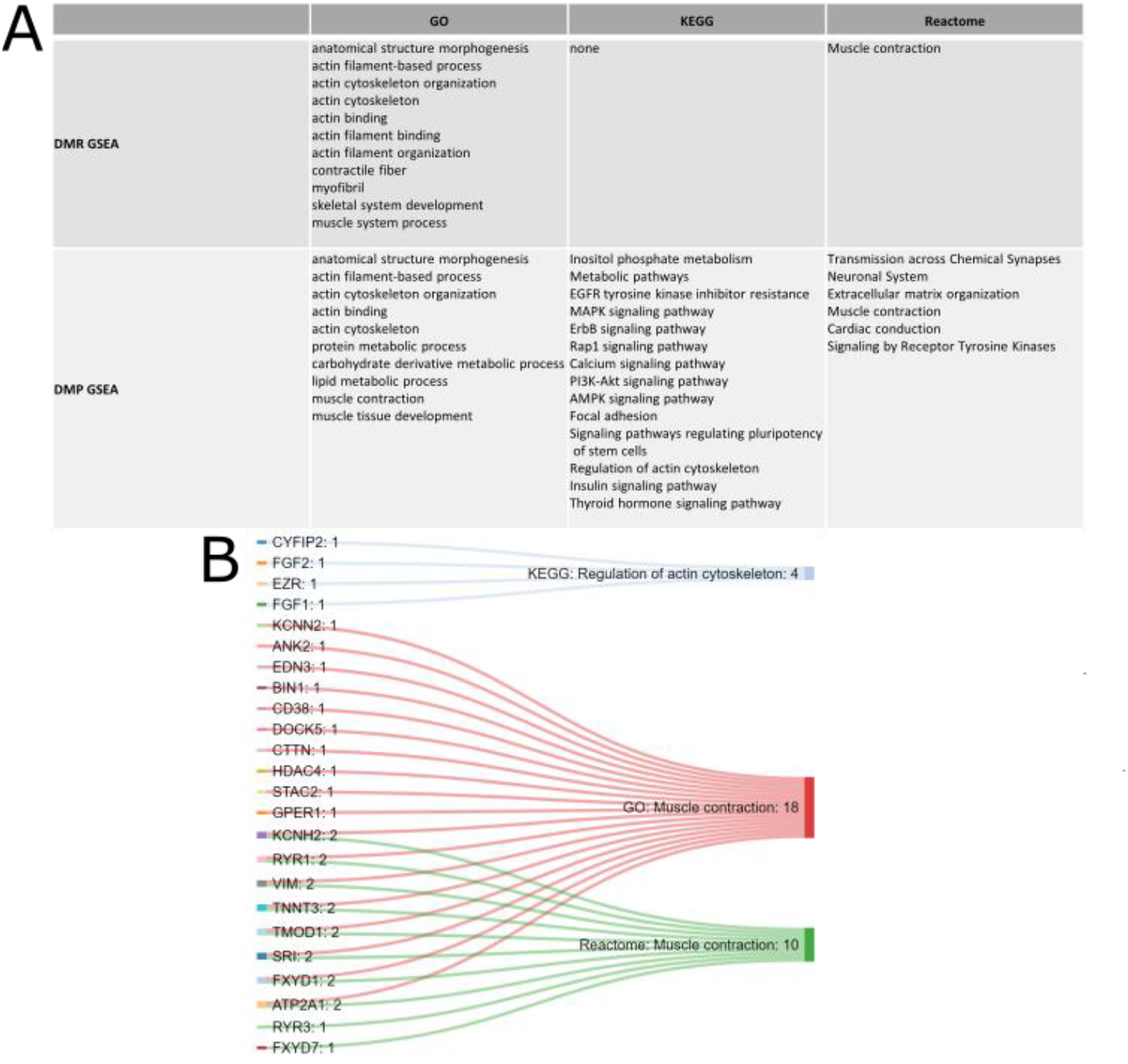
Gene set enrichment analysis of the differentially methylated genes. (**A**) Selected enriched Gene Ontology (GO) terms, Kyoto Encyclopedia of Genes and Genomes (KEGG) pathways, and Reactome pathways from GSEA of DMRs and DMPs. (**B**) Sankey diagram of muscle contraction-related pathways across the three GSEA databases tested and genes within those pathways that were both differentially methylated and expressed (in GTEx and FUSION) between males and females. Numbers next to pathways denote the number of enriched genes in the pathway; numbers next to genes denote the number of pathways (from the ones displayed) that the gene belongs to.

### Validation of GTEx sex-biased genes in the cohorts used for methylation analysis

We sought to validate the sex-biased gene expression obtained from GTEx in a subset of the samples used for methylation analysis since the DMGs and DEGs analyses were obtained from different muscle groups (the DMGs of the current study are from the *vastus lateralis* while the GTEx DEGs are from the *gastrocnemius*). Although both are skeletal muscle tissue from the leg, there may be differences in muscle phenotypes in differing muscle groups [52]. Analysis of RNA sequencing data from the FUSION cohort revealed 3,751 autosomal genes with sex-biased expression (FDR < 0.005). The FDR threshold we chose for the FUSION gene expression data was more stringent than the GTEx local false sign rate threshold (lfsr < 0.05), yet, ~34% of the genes which were both DEGs in GTEx and DMGs were also DEGs in the FUSION cohort, totalling 326 genes (hereinto referred to as ‘overlapping genes’) (**Figure 3A**). Given that both the GTEx and FUSION cohorts include participants of relatively older ages, we sought to confirm the mRNA levels in the younger cohort in the analysis (the Gene SMART) for three genes that displayed sex differences at both the mRNA and DNA methylation levels (*GGT7, FOXO3, and ALDH1A1*) (**Table 2**, Supplementary Table 11, Supplementary Figure 6).

**Table 2.** Gene expression and DNA methylation differences between males and females for three genes across the cohorts used in the analysis.

|  | Gene expression effect size between males and females |  |  | Mean effect size of DMR showing largest effect size (% DNA methylation difference between males and females) |
| --- | --- | --- | --- | --- |
|  | GTEx (MASH posterior effect size) | FUSION (Fold change) | Gene SMART (Fold change) |  |
| *GGT7* | 0.81 | 1.6 | 3.0 | -20.4 |
| *FOXO3* | -0.04 | -0.2 | 3.4 | -1.9 |
| *ALDH1A1* | 0.33 | 0.4 | 2.0 | -10 |

### DNA methylation and gene expression of GGT7, FOXO3 and ALDH1A1 consistently differ between males and females in human skeletal muscle

Three-hundred twenty-six genes exhibited differential methylation in the meta-analysis and differential expression among the GTEx and FUSION cohorts, termed ‘overlapping genes’. Of those genes, we tested three for gene expression levels, *GGT7*, Forkhead Box O3 (*FOXO3*), and Aldehyde Dehydrogenase 1 Family Member A1 (*ALDH1A1*), in the younger cohort included in the DNA methylation analysis (Gene SMART) given the effect that age has on skeletal muscle gene expression [53]. These three genes showed a large effect size in gene expression and DNA methylation, displayed moderate gene expression levels in skeletal muscle relative to other tissues, and/or contained numerous DMPs and DMRs (**Table 2**). The direction of sex-biased expression was consistent for *GGT7* and *ALDH1A1* across GTEx, FUSION, and Gene SMART cohorts (GTEx lfsr < 2.2e^−16^; FUSION FDR= 2.3e^−8^, Gene SMART p-value= 0.03), while the direction was opposite for *FOXO3* (FUSION and GTEx *FOXO3* expression lower in males, Gene SMART *FOXO3* expression higher in males (GTEx lfsr = 0.01; FUSION FDR= 0.001, Gene SMART p-value= 0.002)). As a specific example of the extent of sex differences across the different layers of analysis, *GGT7* displays male-biased expression in skeletal muscle (GTEx lfsr < 2.2e^−16^; FUSION FDR= 1.3e^−45^, Gene SMART p-value= 0.0003) as well as lower methylation in males at DMPs and DMRs annotated to *GGT7* (max DMR: Fisher p-value <0.00^−15^, max beta value effect size=-28.5%, mean beta value effect size=-20.4%) (**Figure 5**).

**Figure 5.**
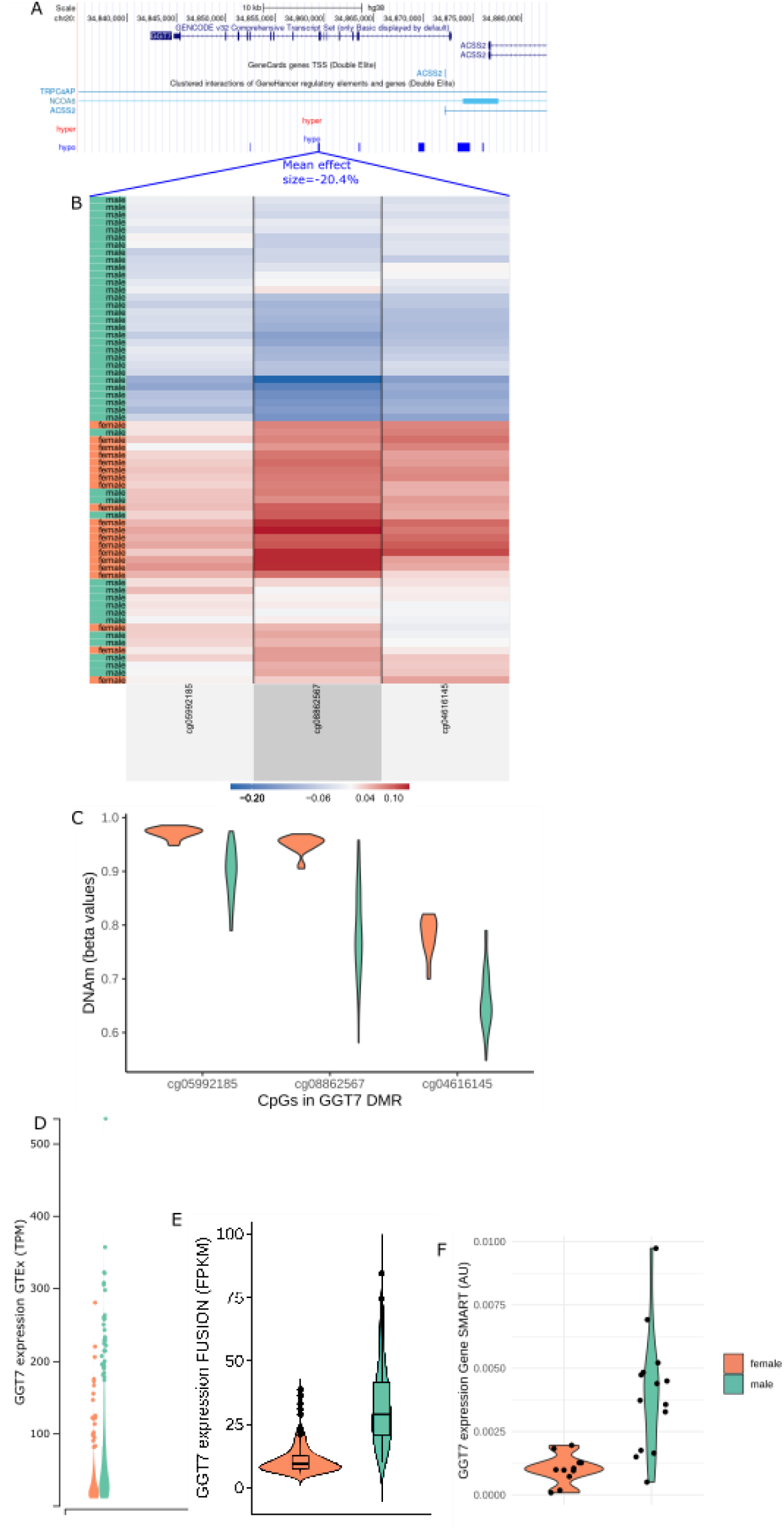
Differential DNA methylation and expression of *GGT7* between males and females. (**A**) UCSC gene track of *GGT7*. From top to bottom: base pair scale in black, GENCODE gene tracks transcript variants in blue, GeneHancer regulatory element annotations in light blue, hyper DMRs tracks in red, hypo DMRs tracks in blue. (**B**) Heatmap of the Gene SMART study (beta values adjusted for all confounders except sex) across the 3 CpGs included in the *GGT7* hypo DMR selected in blue lines and labeled with mean DMR effect size (n=65). Each row represents an individual; green denotes males and orange denotes females; ordered by similarity to other individuals. Each column corresponds to a CpG in the DMR, ordered by genomic location and corresponding to 5C. Blue denotes hypomethylation; tred denotes hypermethylation. (**C**) Distribution of DNA methylation (beta values) in males and females, for the three CpGs in the DMR, matching 5B (n = 65). (**D**) *GGT7* RNAseq expression (TPM-transcripts per million) in males and females of the GTEx (adapted from GTEx portal, n = 803). (**E**) *GGT7* RNAseq expression in the FUSION males and females (FPKM-fragments per kilobase of transcript per million) (n = 274). (**F**) *GGT7* qPCR expression in a subset of Gene SMART males and females (Arbitrary Units; 2^−ΔCt^) (n = 25).

## Discussion

We conducted a large-scale meta-analysis of DNA methylation differences between males and females in skeletal muscle, and integrated them with transcriptomic data. We revealed that males display profound genome-wide hypomethylation compared with females, and that hormone-related TFBSs and muscle fibre type proportions underlie the observed DNA methylation sex differences. We also showed that many sex-biased genes found in GTEx exhibit sex-biased DNA methylation, which was partially confirmed in the FUSION cohort. We then validated the gene expression (qPCR) levels of three genes with large DNA methylation and expression differences between the sexes across cohorts, and confirmed the higher gene expression in males of *GGT7* and *ALDH1A1*. Finally, we showed that the DMGs are overwhelmingly involved in muscle contraction, as well as substrate metabolism and anatomical structure-related pathways.

In the present study, the overwhelming majority (94%) of the DMPs were hypomethylated in males. Interestingly, global autosomal hypomethylation in males has been observed in various other tissues [54], including blood [55, 56] and pancreatic islets [20]. We aimed to uncover some of the biological mechanisms at the root of these epigenetic sex-differences by assessing the contributions of fibre type proportions and sex hormone levels to the observed DNA methylation sex differences. We hypothesised that differences in fibre type proportions between sexes may partly explain our findings [56–58], as studies report that type I fibres are hypermethylated compared with type II fibres [40], and as females tend to have a higher proportion of type I fibres than males [18]. Consistent with this, we observed that females had higher proportions of type I muscle fibres than males and that type I fibre content was mostly associated with DNA hypermethylation. Importantly, 16% of the loci exhibiting sex-biased DNA methylation were also associated with fibre type proportions. This suggests that at those CpGs, differences in DNA methylation between the sexes is due to the inherent sex differences in fibre type proportions. Nonetheless, the vast majority of the loci that exhibit sex-biased DNA methylation (84%; 48,008 CpGs) differ regardless of the sex differences in fibre type proportions. A recent study on the FUSION cohort, adjusted for fibre type proportions and found that it explains a substantial portion of the variability in DNA methylation for many metabolic phenotypes of interest [59]. Skeletal muscle DNA methylation analyses are performed on whole muscle due to the cost and technical limitations of isolating muscle cell types. Differing non-muscle cell types may be present in a muscle biopsy sample and it is currently unknown how much of the muscle DNA methylation profile may actually be representing other cell types [60–62]. Bioinformatics deconvolution methods have not yet been developed for bulk skeletal muscle DNA methylation. Considering that the DNA methylation differences between cell types are large [63], future studies should aim at determining DNA methylation patterns of the different muscle fibre and cell types so that bulk muscle DNA methylation data can be adjusted for the appropriate cell and fibre proportions.

None of the circulating sex hormones were associated with differential methylation across all CpGs, nor across the sex-DMPs in males or females. However, the range of each hormone within each sex may not be large enough to draw out the effect of varying levels of testosterone on the methylome. In the current study, hormone levels were measured from blood while DNA methylation was measured from skeletal muscle. DNA methylation patterns are highly tissue-specific [64, 65] and sex hormone levels in the circulation are not necessarily correlated with those intramuscularly. Moreover, intramuscular, and not circulating, sex hormone levels may be correlated with muscular function [66, 67]. A recent review emphasizes the importance of measuring intramuscular sex hormone levels when assessing muscle-related properties in females [68].

The enrichment of hormone-related TFBS among the sex-DMPs suggests that lifelong exposure to differing hormone levels significantly contributes to the observed sex differences in skeletal muscle DNA methylation. In Unibind, ChIP-seq data in skeletal muscle was limited to one TF (CTCF), so the enrichment of TFBSs among sex DMPs may have limited functional significance in skeletal muscle. Nonetheless, many of the TFs that showed strong enrichment in the present study, such as *AR* [69, 70], *ESR1* [70], and *SMAD3* [71] are expressed in skeletal muscle and have important roles in muscle phenotype. Two recent studies leveraging the GTEx database identified sex differences in TF targeting patterns across several human tissues, including skeletal muscle, which contribute to sex-biased gene regulatory networks [14] and gene expression [7]. Differences in sex hormone levels between developing males and females are already evident *in utero* [72], making it challenging to design an experiment in humans that disentangles the effect of long-term hormonal exposure from biological sex, and other related factors, on cell function. Studies have utilized menopausal females [73] and transgender people [74] receiving hormone replacement therapy (HRT) to investigate the influence of long term exposure to sex hormones on various phenotypes and risk of diseases. For example, HRT for one postmenopausal monozygotic twin and not the other has positive effects on regulation of muscle contraction and myonuclei organization, suggesting that oestrogen has direct effects on muscle function [75]. Nevertheless, uncovering the genomic regions that display sex-differential methylation as well as contain hormone-responsive TFBSs, provides insight on which genomic regions, hormones, and TFs are discerning male and female skeletal muscle.

The enrichment for muscle contraction-related pathways among the DMGs across GO, KEGG, and Reactome suggests that the sex-differential DNA methylation has functional relevance in skeletal muscle function. Furthermore, enrichment of substrate metabolism pathways among the DMGs suggests that the observed sex differences in carbohydrate, lipid, and protein metabolism [16, 76] may have a molecular, more specifically an epigenetic, basis. This is corroborated with results from transcriptomic studies, which report that skeletal muscle female-biased genes are enriched for pathways involved in fatty acid metabolism while male-biased genes are enriched for pathways involved in protein catabolism (Lindholm et al., 2014a).

Although not well-understood, the sex chromosome complement may also influence autosomal DNA methylation patterns. In cultured fibroblasts, the presence of Sex-determining Region Y (*SRY*) is associated with lower autosomal methylation levels [77–79]. Additionally, a higher number of the X chromosomes, in the absence of *SRY*, leads to increased methylation levels at a specific sex-differentially methylated autosomal region [79]. This could be attributed to allele dosage compensation, a female-specific process that silences one of the X chromosomes in a cell [80, 81]. Approximately one-third of genes ‘escape’ inactivation, remain transcriptionally active in XX cells, [81–83], and have been suggested to affect autosomal DNA methylation via their histone marks [79, 84]. Moreover, females with Turner syndrome (partially/fully missing one X) and monosomy X have lower global methylation than XX females, but higher than XY males [85, 86]. The effect of sex chromosome complement on autosomal DNA methylation in skeletal muscle has yet to be explored. Finally, genetic variants (copy number variants (CNVs) and single nucleotide polymorphisms (SNPs)) may affect DNA methylation status at specific loci [87], termed methylation-quantitative trait loci (meQTL). However, with the exception of a female bias for large, rare CNVs, there are no large sex differences in autosomal SNP minor allele frequencies [3].

The relationship between DNA methylation and gene expression is complex; DNA methylation at promoters, enhancers, and 1^st^ exons is generally believed to enhance gene silencing, while DNA methylation at gene bodies can sometimes be associated with increased gene expression (Rauch et al., 2009, Lister et al., 2009, Ball et al., 2009, Aran et al., 2011, Jones, 2012). Using a permutation test, we showed that DNA methylation differences between the sexes at promoters and enhancers were more often associated with lower gene expression than would be expected by chance alone. DNA methylation differences between the sexes were also particularly prominent in chromatin states that are known differ between males and females. This suggests that DNA methylation differences between males and females reflect alterations in chromatin activity, and differential epigenetic states and expression are likely functionally connected. In line with this, chromatin states that differ between the sexes have been shown to be enriched for sex-biased genes across various tissues, including skeletal muscle [46]. However, it is not yet possible to assess whether the relationship reflects correlation or direct causality. There is still debate around whether epigenomic features drive regulatory processes or are merely a consequence of transcription factor binding [46].

We identified 326 genes with consistent differential skeletal muscle DNA methylation and expression across 1,172 individuals altogether (369 individuals from three cohorts for DNA methylation and 1,077 individuals from two cohorts for gene expression). Although we found profound global DNA hypomethylation in males, of the overlapping genes there were equivalent numbers of genes over- and under-expressed in males compared with females for both GTEx and FUSION. Indeed, hypermethylation is not always associated with decreased gene expression [88]. The substantial overlap between differentially methylated genes and differentially expressed genes highlights many genes that may be of interest for their roles in muscle-related processes. We focused on three of these genes that displayed a large DNA methylation difference between males and females, are highly expressed in skeletal muscle, or play a role in skeletal muscle function: *HDAC4* given its role in neurogenic muscle atrophy [89, 90] and in the response to exercise [91]; *DEPTOR* given its role in muscle glucose sensing which in turn augments insulin action [92]; *GRB10* given that it is imprinted and has been shown to change in methylation with exercise/training [93]; *FOXO3* for its role in ageing, longevity, and regulating the cell cycle [94]; *ALDH1A1* for its role in aldehyde oxidation and because sex differences in skeletal muscle mRNA levels have been reported, suggesting that males might be able to metabolize aldehydes (i.e. alcohol) more efficiently than females [15]; and *GGT7* for its role in antioxidant activity [95]. Of the three genes which were validated across GTEx, FUSION, and Gene SMART, two of them showed consistently higher male expression levels (*GGT7, ALDH1A1*) while one showed opposite sex-biased expression (*FOXO3*) in the young versus the old cohorts. *FOXO3* expression was lower in males in the older cohorts (GTEx and FUSION), and higher in males in the younger cohort (Gene SMART). Other studies have shown that males have higher *FOXO3* expression in young skeletal muscle [96] and that elderly females have higher skeletal muscle *FOXO3* expression than younger females [97]. While *FOXO3* skeletal muscle gene expression differs between males and females, it seems that the direction is opposite in young and old individuals, which emphasizes the caution that should be used when interpreting sex differences across a large age range of individuals. The promoter, 1^st^ exon, and gene body of *GGT7* were hypomethylated in males and males had higher *GGT7* expression. *GGT7* is highly expressed in skeletal muscle and metabolises glutathione, which is a ubiquitous “master antioxidant” that contributes to cellular homeostasis. Efficient glutathione synthesis and high levels of glutathione-dependent enzymes are characteristic features of healthy skeletal muscle and are also involved in muscle contraction regulation [98].

In conclusion, we showed that the DNA methylation of hundreds of genes differs between male and female human skeletal muscle. We uncovered important biological factors underlying sex-specific skeletal muscle DNA methylation. Integration of the DNA methylome and transcriptome, as well as gene expression validation, identify sex-specific genes associated with muscle metabolism and function. Uncovering the molecular basis of sex differences across different tissues will aid in the characterization of muscle phenotypes in health and disease. The effects of other upstream drivers on sex differences in the muscle methylome, such as non-muscle cell type, the XY chromosomes, and genetic variants still need to be explored. Molecular mechanisms that display sex differences in skeletal muscle may help uncover novel targets for therapeutic interventions.

## Methods

### Datasets

We conducted a meta-analysis of three independent epigenome-wide association studies (EWAS) of sex including the Gene Skeletal Muscle Adaptive Response to Training (SMART) study from our lab [99], the Finland-United States Investigation of NIDDM Genetics (FUSION) study from the dbGAP repository (phs000867.v1.p1) [59], and the GSE38291 dataset from the Gene Expression Omnibus (GEO) platform [100]. Detailed participant characteristics, study design, muscle collection, data preprocessing, and data analysis specifications for each study are in Supplementary table 1. Briefly, all studies performed biopsies on the *vastus lateralis* muscle, all participants were of Caucasian descent (except 1 individual of mixed Caucasian/aboriginal decent), and included either healthy or healthy and T2D individuals aged 18-80 years. The Gene SMART study was approved by the Victoria University human ethics committee (HRE13-223) and written informed consent was obtained from each participant. NIH has approved our request [#96795-2] for the dataset general research use in the FUSION tissue biopsy study.

### DNA Extraction and Methylation Method-Gene SMART study samples

Genomic DNA was extracted from the samples using the AllPrep DNA/RNA MiniKit (Qiagen, 80204) following the user manual guidelines. Global DNA methylation profiling was generated with the Infinium MethylationEPIC BeadChip Kit (Queensland University of Technology and Diagenode, Austria). The first batch contained only males and were randomized for timepoint and age. The second batch contained males and females and samples were scrambled on the chips to ensure randomness when correcting for batch effect (i.e. old and young males and females included on each chip). The genome-wide DNA methylation pattern was analysed with the Infinium MethylationEPIC BeadChip array.

### Bioinformatics and statistical analysis of DNA Methylation

#### PREPROCESSING

The pre-processing of DNA methylation data was performed according to the bioinformatics pipeline developed for the Bioconductor project [101]. Raw methylation data were pre-processed, filtered and normalized across samples. Probes that had a detection p-value of > 0.01, located on X and Y chromosomes or cross-hybridizing, or related to a SNP frequent in European populations, were removed. It is important to note that the list of cross-hybridizing probes was supplied manually [102] as the list supplied to the *ChAMP* package was outdated. Specifically, there are thousands of probes in the Illumina microarrays that cross-hybridize with the X-chromosome and may lead to false discovery of autosomal sex-associated DNA methylation [103]. The BMIQ algorithm was used to correct for the Infinium type I and type II probe bias. β-values were corrected for both batch and position in the batch using *ComBat* [104].

#### STATISTICAL ANALYSIS

We adjusted each EWAS for bias and inflation using the empirical null distribution as implemented in *BACON* [1]. Inflation and bias in EWAS are caused by unmeasured technical and biological confounding, such as population substructure, batch effects, and cellular heterogeneity [105]. The inflation factor is higher when the expected number of true associations is high (as it is for age); it is also greater for studies with higher statistical power [1]. The results were consistent with the inflation factors and biases reported in an EWAS of age in blood [1]. Results from the independent EWAS were combined using an inverse variance weighted meta-analysis with METAL [106]. We used METAL since it does not require all DNA methylation datasets to include every CpG site on the HumanMethylation arrays. For robustness, we only included CpGs present in at least 2 of the 3 cohorts (633,645 CpGs). We used a fixed effects (as opposed to random effects) meta-analysis, assuming one true effect size of sex on DNA methylation, which is shared by all the included studies. Nevertheless, Cochran’s Q-test for heterogeneity was performed to test whether effect sizes were homogeneous between studies (a heterogeneity index (I2) > 50% reflects heterogeneity between studies).

To identify DMPs, we used linear models as implemented in the *limma* package in R [107], using the participants’ ID as a blocking variable to account for the repeated measures design (for twin (GSE38291) and duplicate samples (Gene SMART), using DuplicateCorrelation). The main sources of variability in methylation varied depending on the cohort and were adjusted for in the linear model accordingly. For the Gene SMART study, we adjusted the linear models for age, batch (2017 vs 2019), sex, timepoint and the interaction of sex and timepoint (before and after four weeks of high-intensity interval training). For the FUSION study, we adjusted the linear models for age, sex, BMI, smoking status, and OGTT status. For the GSE38291 study, we adjusted the linear models for age, sex, and diabetes status. All results were adjusted for multiple testing using the Benjamini and Hochberg correction and all CpGs showing an FDR < 0.005 were considered significant [109]. DMRs were identified using the *DMRcate* package [110]. DMRs with Stouffer, Fisher, and harmonic mean of the individual component FDRs (HMFDR) statistics < 0.005 were deemed significant. Effect sizes are reported as mean differences in DNA methylation (%) between the sexes.

To assess whether fibre type proportions differ between males and females in the Gene SMART and FUSION cohorts we used a beta regression model [111] using the *betareg* package in R. We then included type I fibre ratio as a covariate in the linear models. To investigate whether circulating hormones is associated with DNA methylation of the sex-DMPs, we included each sex hormone as a covariate in a linear model, separately. One male from the Gene SMART cohort had missing circulating hormone values; and one two females from the Gene SMART cohort had missing type I fibre proportions. For these three individuals missing values were imputed with the *mice* package in R [112].

We integrated a comprehensive annotation of Illumina HumanMethylation arrays [113] with chromatin states from the Roadmap Epigenomics Project [45] and the latest GeneHancer information [114]. DMPs that were annotated to two differing chromatin states were removed for simplicity and because there were very few such DMPs. GSEA on KEGG and GO databases was performed on DMRs and DMPs using the *goregion* and *gometh* (*gsameth* for Reactome) functions in the *missMethyl* R package [115] [116].

Enrichment of TFBSs among the identified DMRs was performed using the enrichment analysis tool in http://unibind.uio.no/ which utilizes the *runLOLA* function of the R package *LOLA* [48].

### Fibre types: meta analysis and derivation from immunohistochemistry and RNA-seq

Myosin heavy chain is currently the best available marker for fibre typing [117]. Gene SMART muscle sections were frozen in optimum-cutting temperature (OCT) medium by holding the sample with OCT in liquid-nitrogen cooled Isopentane until frozen. Samples were stored in −80°C until they were sectioned at 8μM with a cryostat. The IHC protocol was performed as is described elsewhere [118]. Briefly, sections were blocked in 4% goat serum (Invitrogen). Primary antibodies BA-F8 (MHCI), BF-35 (MHCIIA), and 6H1 (MHCIIX) were purchased from DSHB, Iowa. Secondary antibodies goat anti-mouse IgG2b 350, goat anti-mouse IgG1 488, and goat anti-mouse IgM 555 were purchased from Invitrogen. Some samples were fixed in paraformaldehyde for other analyses and for those samples an antigen-retrieval protocol consisting of a 10 min incubation at 50° C of Proteinase K diluted in milliQ (1:1000) and subsequent 1 min washes was performed before the IHC. Imaging was performed on the Olympus BX51.

To determine type I fibre proportions in the FUSION cohort we followed the validated method as performed by the original study on the FUSION cohort [59]. Briefly, we derived type I fibre proportions from the RNA-seq expression data (TPMs) for type I (MYH7), type IIA (MYH2), and type IIX (MYH1). We calculated the ratio of MYH7 out of the total. We then included this ratio in the +FT linear model.

To determine whether the inherent sex differences in fibre type proportions underlie the sex differences in DNA methylation we separated the males and females of the Gene SMART and FUSION cohorts and performed a meta-analysis on the four groups (FUSION females, FUSION males, Gene SMART females, Gene SMART males). Given that females displayed significantly higher type I fibre proportions than males in both cohorts, we could not simply include type I fibre content in a linear model performed on a mixed sex cohort as two issues would arise: i) collinearity of fibre type with sex ii) differences in fibre type proportions may be a downstream effect of sex. Dividing the cohorts by sex, conducting a meta-analysis, and selecting the sex-biased DMPs and performing an FDR adjustment among those cites allowed us to address whether fibre type proportion is associated with DNA methylation at sex-biased DNA methylation loci. The fibre type meta-analysis was performed with the same methodology of the sex meta-analysis as described above.

### Controlling for the female menstrual cycle

Various contraceptives have different dosage, administration patterns, and different hormone combinations causing variability in metabolism and gene expression [119], therefore only females not taking any form of hormonal contraceptives were recruited for the Gene SMART study. Furthermore, to minimise the effect of fluctuating hormone levels, females were required to have a regular menstrual cycle (27-35 days), and all samples were aimed to be collected during the early follicular phase (day 1-day 8 of cycle), with few exceptions due to logistics. Oestrogen, progesterone, follicle stimulating hormone (FSH), and luteinizing hormone (LH) were measured in blood serum. Given the intricate fluctuations of the ovarian hormones, these four hormones were combined into a principal component analysis and the first two PCs, which explained the majority of the variability, were each added into the linear model.

### Blood serum hormones

The hormone assays were completed in the accredited pathology laboratory at Monash Health, Australia. Estradiol (E2) and Progesterone assays are competitive binding immunoenzymatic assays performed on the Unicel DXI 800 system (Beckman Coulter). FSH assay is based on Microparticle Enzyme Immunoassay (MEIA) and is carried out on the Unicel DXI 800 system (Beckman Coulter). The LH and sex-hormone binding globulin (SHBG) assays were performed using a sequential two-step immunoenzymatic (“sandwich”) assay carried out on a Unicel DXI 800 (Beckman Coulter). Testosterone was measured using the HPLC–tandem mass spectrometry method using a liquid sample extraction (AB Sciex Triple Quad 5500 liquid chromatography–tandem mass spectrometry). Free androgen index was calculated as (total testosterone × 100)/SHBG. Free testosterone (fT) was calculated by the Södergard fT calculation (36).

### Integration of DNA Methylation and Gene Expression

The Genotype-Tissue Expression (GTEx) Project sex-biased data was downloaded from the GTEx Portal on 08/26/2020 and filtered for skeletal muscle samples. The enrichment of DMG for GTEx DEGs was done by supplying the list of sex-biased genes to the *gsameth* function in the *missMethyl* R package [115, 116], which performs a hypergeometric test, taking into account biases due to the number of CpG sites per gene and the number of genes per probe on the EPIC array. Therefore, caution should be taken when interpreting the number of DMPs reported per DMG. The analysis for direction of correlation between DNA methylation and gene expression was performed by randomly shuffling DNA methylation effect sizes and performing 10,000 permutations to assess how often a negative correlation occurs. This analysis was performed for both GTEx and FUSION transcriptome data and yielded similar results; data presented reflect results from the integration of differential methylation with differential GTEx expression. Significance reported for GTEx sex-biased genes is represented as the local false sign rate (lfsr) which is analogous to FDR [120]. GTEx effect sizes are represented as mash posterior effect sizes [120], in which positive values indicate male-biased genes and negative values indicate female-biased genes. FUSION and Gene SMART gene expression significance statistics are represented as FDR and p-value, respectively, and effect sizes as fold changes for both cohorts.

### Validation of top genes with qPCR

Skeletal muscle previously stored at −80°C was lysed with the RLT buffer Plus buffer (Qiagen) and beta-mercaptoethanol using the TissueLyser II (Qiagen, Australia). DNA was extracted using the AllPrep DNA/RNA Mini Kit following the manufacturer guidelines (Qiagen, Australia). RNA yield and purity were assessed using the spectrophotometer (NanoDrop One, Thermofisher). RNA was reverse transcribed to cDNA using a commercially available iScript Reverse Transcriptase supermix (cat #1708841) and C1000 Touch Thermal Cycler (Bio-Rad, Hercules, CA, USA). Complementary DNA samples were stored at −20°C until further analysis. Quantitative real-time PCR was performed using SYBR Green Supermix (Bio-Rad, Hercules, CA) and gene-specific primers (listed in Supplementary table 11). Primers were either adapted from existing literature or designed using Primer-BLAST (http://www.ncbi.nlm.nih.gov/tools/primer-blast/) to include all splice variants, and were purchased from Integrated DNA Tehcnologies. Ten microliter reactions comprised of SYBR, and optimised concentrations of forward and reverse primers (Supplementary table 11 for primer conditions), nuclease free water and 8 ng of cDNA were run in triplicate using an automated pipetting system (epMotion M5073, Eppendorf, Hamburg, Germany), with no-template negative controls on a 384-well plate in a thermo-cycler (QuantStudio 12K Flex System, ThermoFisher Scientific, Australia). Gene expression was normalized to the geometric mean expression of the two most stable housekeeping genes, as determined by Ref finder, TATAA-box binding protein (TPB), and 18s rRNA, which did not differ between sexes (Supplementary table 11). Data are presented as the fold change in males compared to females, using 2^−ΔΔCT^.

## Acknowledgements

We would like to thank Dr Alba Moreno-Asso from Victoria University for her assistance with the qPCR analysis and Dr Andrew Garnham for his expertise performing the muscle biopsies. The Genotype-Tissue Expression (GTEx) Project was supported by the Common Fund of the Office of the Director of the National Institutes of Health, and by NCI, NHGRI, NHLBI, NIDA, NIMH, and NINDS. The data used for the analyses described in this manuscript were obtained from the GTEx Portal on 08/26/2020.

## Contributions

Conceptualization: S.L., S.V., and N.E. Methodology: S.L., and S.V. Investigation: S.L., S.V. Formal analysis: S.L. Resources: S.L., M.J., D.H., J.A.R., N.R.H., L.M.H., L.R.G., K.J.A., and N.E. Writing—Original draft: S.L., S.V., D.H., and N.E. Writing—Review & editing: S.L., S.V., D.H., S.Lamon, N.R.H., L.M.H., K.J.A., and N.E. Funding acquisition: S.V., L.R.G., and N.E.

## Availability of Data and Materials

The dataset generated and analysed during the current study are available in the GEO repository, https://www.ncbi.nlm.nih.gov/geo/query/acc.cgi?acc=GSE171140.

## Consent for publication

Not applicable.

## Supplementary Figures

**Supplementary Figure 1.**
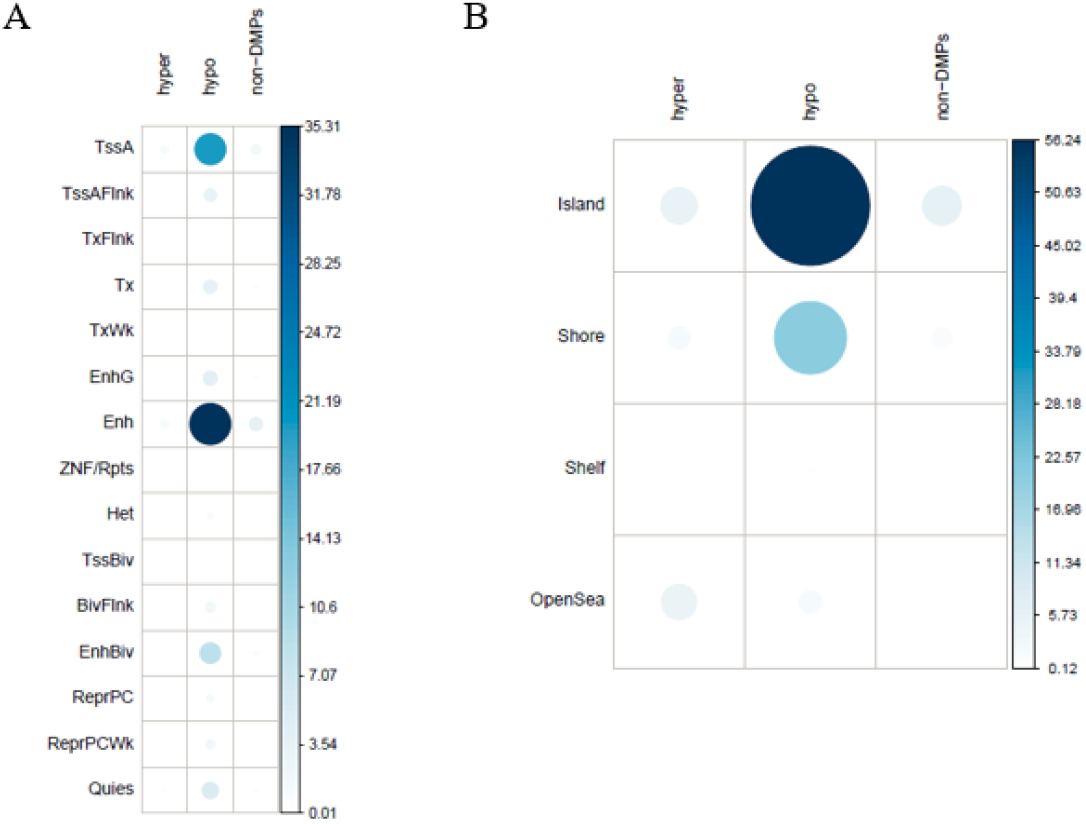
Correlation plots of chi2 tests of genomic locations. (**A**) Correlation plot of the percent contributions to the chi2 test for chromatin states in hyper-, hypo-, and non-DMPs. This plot is using the male chromatin state annotation in skeletal muscle but the female chromatin state annotation revealed equivalent results. Darker blue indicates a greater contribution to the significant relationship between DMP status and chromatin state. (**B**) Correlation plot of the percent contributions to the chi2 test for CGI status of hyper-, hypo-, and non-DMPs. Darker blue indicates a greater contribution to the significant relationship between DMP status and CGI status.

**Supplementary Figure 2.**
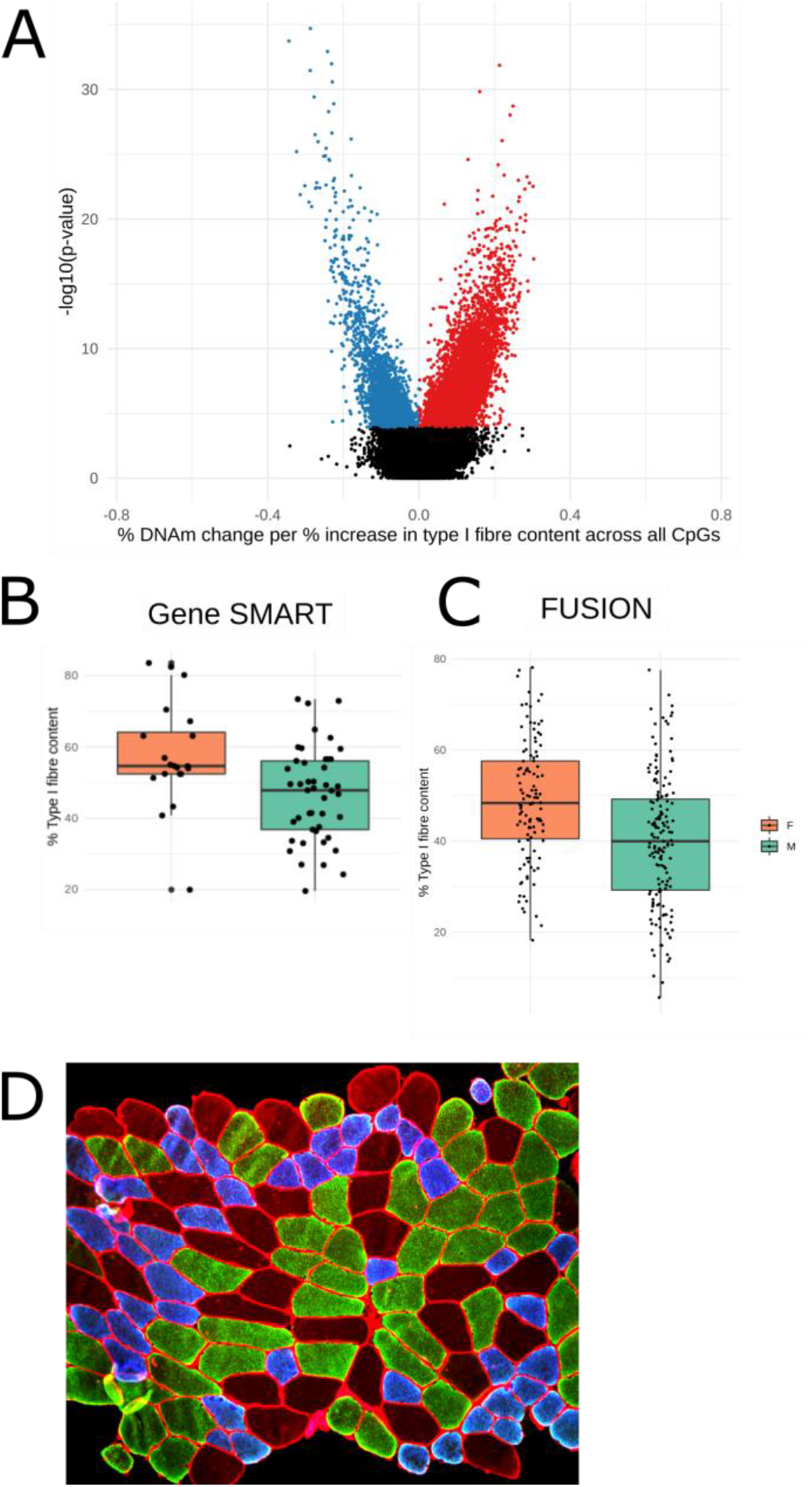
Fibre type proportion analysis. **(A)** Differentially methylated positions (DMPs) with type I fibre proportion across all CpGs conducted with a meta-analysis of males and females, separately, from the Gene SMART and FUSION cohorts. Volcano plot of DNA methylation changes per percent increase in type I fibre content (expressed at percentage of beta value). Each point represents a tested CpG (665,904 in total) and those that appear in color are DMPs at a false discovery rate < 0.005; red DMPs are hypermethylated in type I fibres; blue DMPs are hypomethylated in type I fibres. The x-axis represents the amount of DNA methylation difference with increasing type I fibre content and the y-axis represents statistical significance (higher = more significant). **(B)** Percent of type I fibres in Gene SMART female and male muscle samples as determined by immunohistochemistry; p-value = 0.0001; 59% in females versus 47% in males. **(C)** Percent of type I fibres in FUSION female and male muscle samples as determined by RNA-seq; p-value = 2.06 x 10-7; 48.6% in females versus 39.9% in males. **(D)** Immunohistochemistry (IHC) myosin heavy chain staining of skeletal muscle section. Example of cross-sectional fibres of one participant. Blue fibres indicate type I, green indicate type IIa, and red indicate type IIx; cell membrane staining in red. A minimum of 100 fibres counted per person for approximation of fibre type proportions.

**Supplementary Figure 3.**
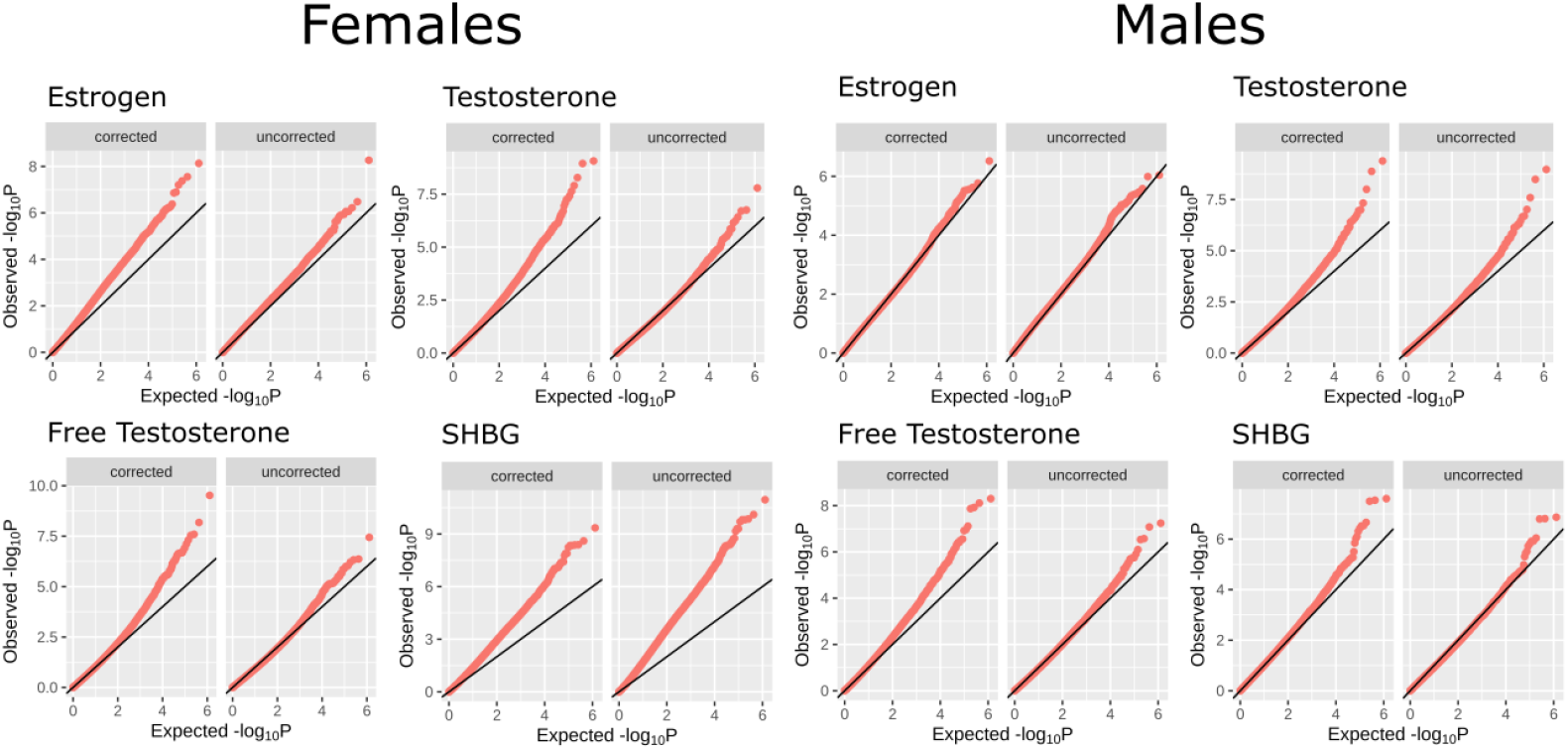
Quantile-quantile (QQ) plots of the analysis of DNA methylation and circulating sex hormones in males and females from the Gene SMART study. Each Epigenome-wide association study (EWAS) was corrected for with *BACON* in R [1], as labelled “corrected”. The uncorrected QQ plots correspond to the unadjusted p-values from the EWAS.

**Supplementary Figure 4.**
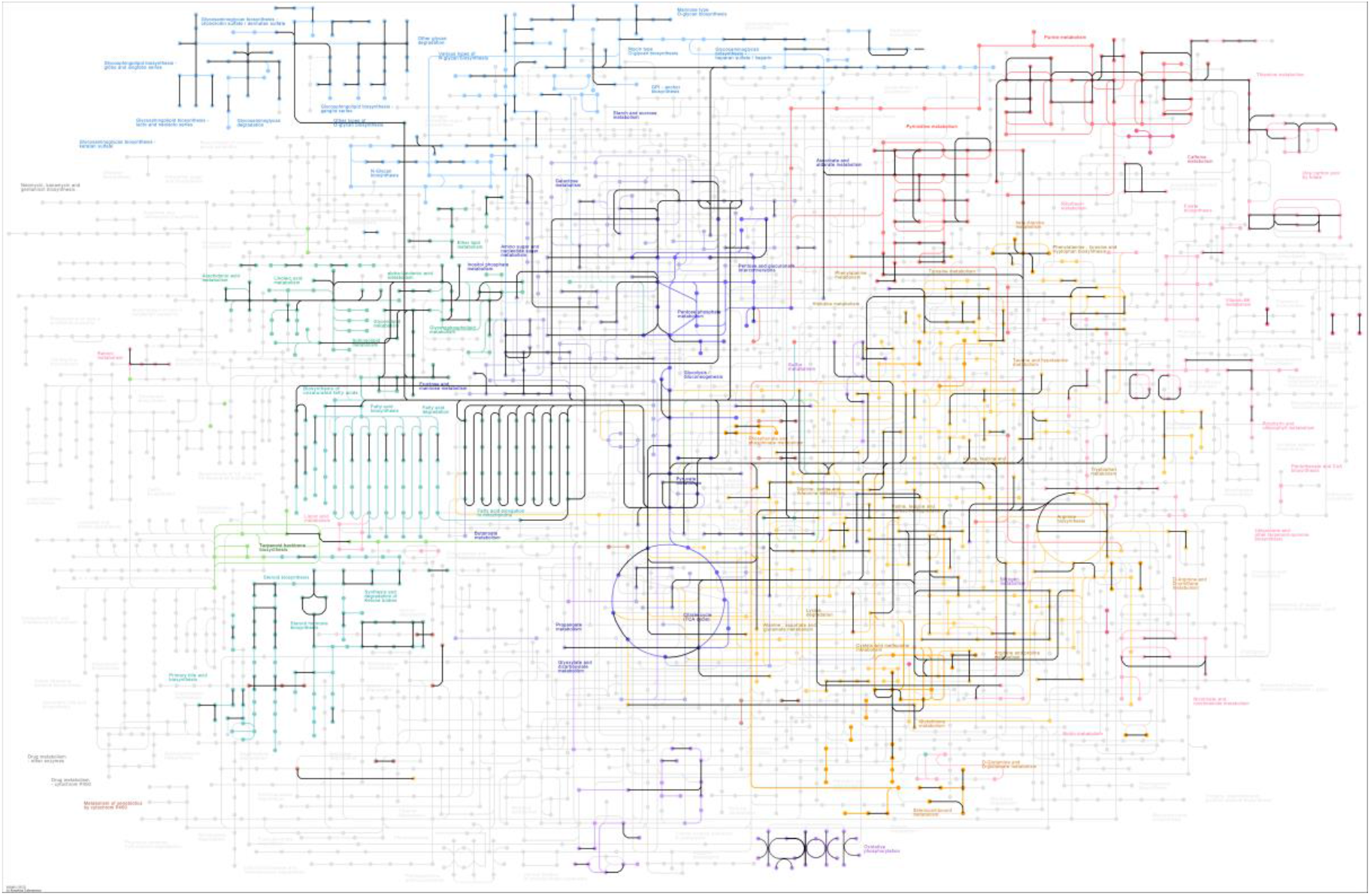
Differentially methylated genes within the KEGG metabolic pathways map (hsa01100). Genes and components of the pathways which display differential methylation between males and females are outlined in black, KEGG GSEA using DMPs.

**Supplementary Figure 5.**
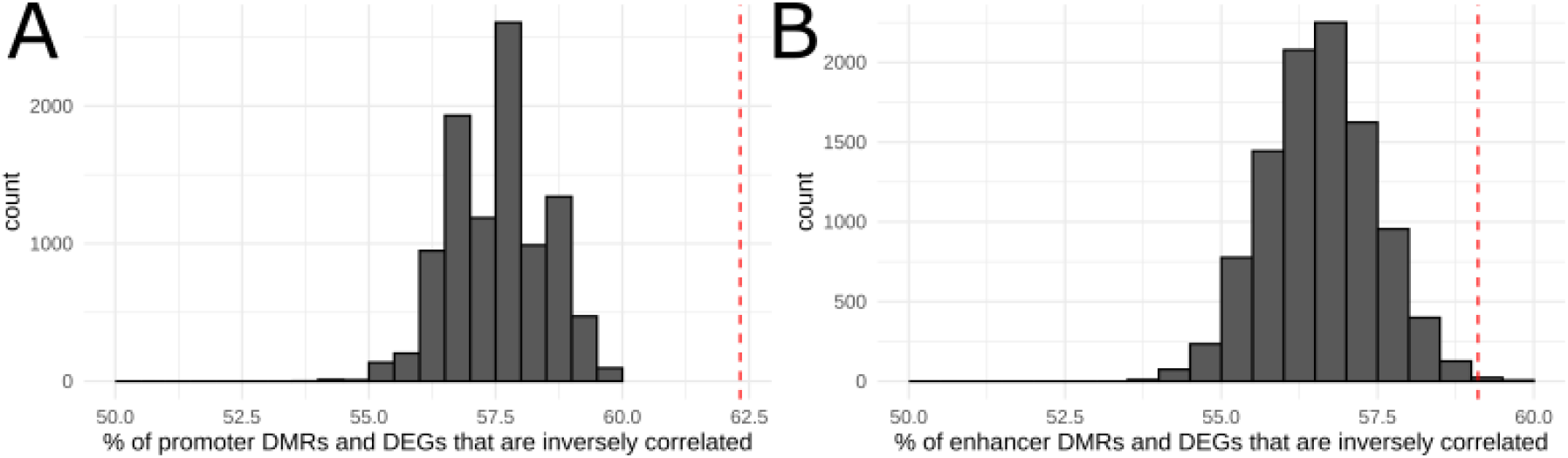
Distribution of the 10,000 random permutations for a negative correlation between DNA methylation and gene expression. (**A**) Histogram of 10,000 random permutations of DMRs annotated to promoter regions and correlation with GTEx gene expression. Effect sizes of DMRs were randomly shuffled and the resulting correlation with gene expression is plotted. Red dashed line indicated the real percentage of promoter DMRs that are negatively correlated with gene expression. (**B**) Histogram of 10,000 random permutations of DMRs annotated to enhancer regions and correlation with GTEx gene expression. Effect sizes of DMRs were randomly shuffled and the resulting correlation with gene expression is plotted. Red dashed line indicated the real percentage of enhancer DMRs that are negatively correlated with gene expression.

**Supplementary Figure 6.**
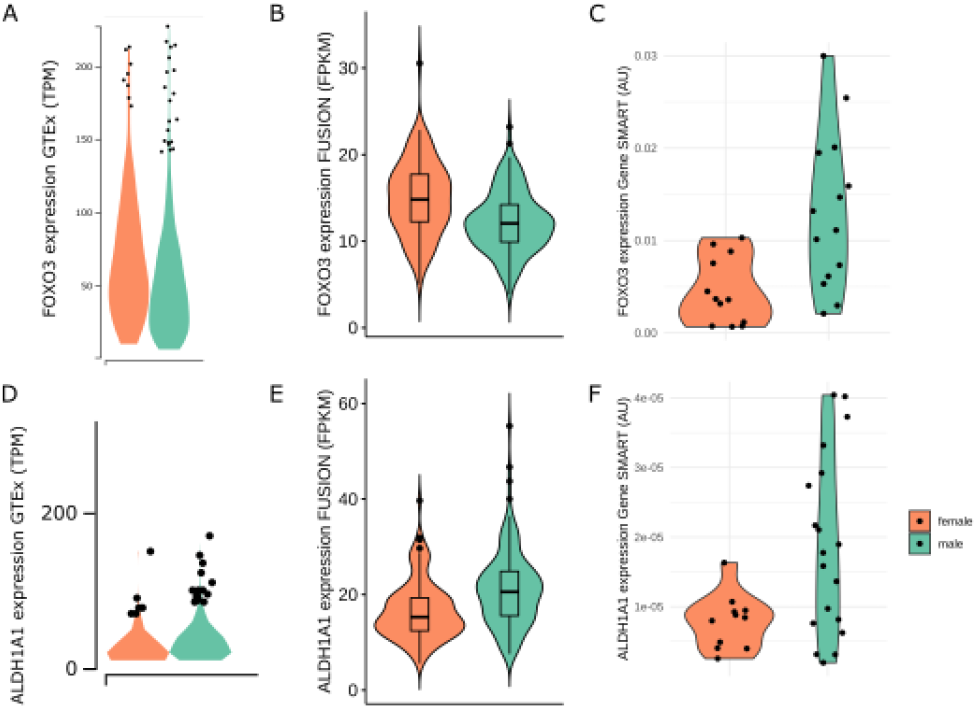
Gene expression for FOXO3 and ALDH1A1 validated in 3 cohorts. Distribution of FOXO3 expression in males and females from (**A**) the GTEx (RNA-seq) (**B**) the FUSION cohort (RNA-seq) (**C**) the Gene SMART cohort (qPCR). Distribution of ALDH1A1 expression in males and females from (**D**) the GTEx (RNA-seq) (**E**) the FUSION cohort (RNA-seq) (**F**) the Gene SMART cohort (qPCR).

**Supplementary Figure 7.**
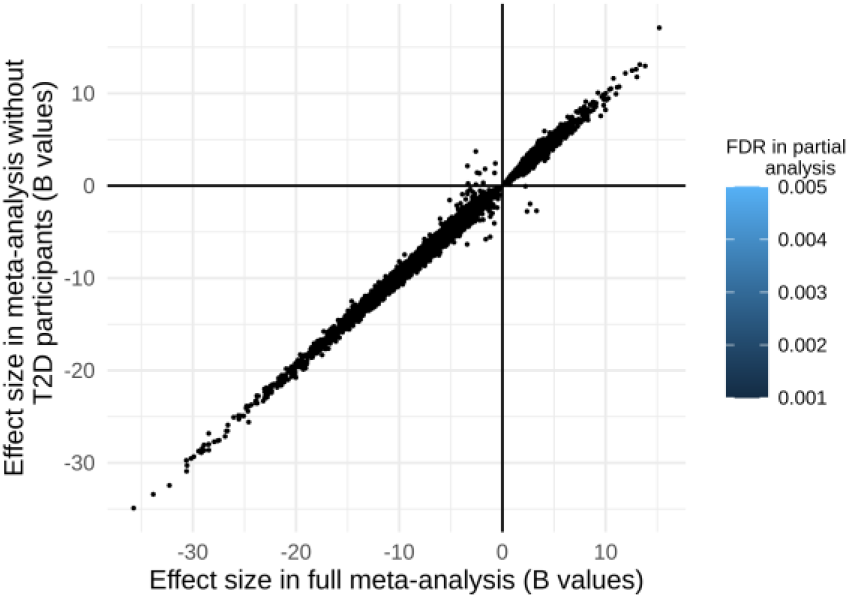
Comparison of results from the full meta-analysis and from a meta-analysis excluding T2D participants in FUSION. Each point is one of the 56,813 differentially methylated positions (DMPs) discovered in the full meta-analysis. To compare results from the full and partial meta-analysis we plotted the effect size (B value percentages) in the full meta-analysis (x-axis) against the effect size of the partial meta-analysis (y-axis). To show whether DMPs remained significant in the partial meta-analysis, we coloured points according to the false discovery rate (FDR) in the partial meta-analysis.

## Supplementary tables legend

***Supplementary table 1-*** Study descriptions. Description of participants, study design, muscle collection, and data preprocessing/analysis.

***Supplementary table 2-*** DMPs. Differentially methylated positions between males and females in the meta-analysis FDR<0.005. Corresponding chromosome, genomic location, annotated genes, male and female chromatin states from the Roadmaps Epigenomics Project, and genes annotated by GeneHancer. Positive effect size indicates higher DNA methylation in males compared to females.

***Supplementary table 3-*** DMRs. Differentially methylated regions between males and females in the meta-analysis Stouffer, HMFDR, and Fisher <0.005. Corresponding chromosome, genomic location, width of DMR, number of CpGs in DMR, statistics (Stouffer, Harmonic mean false discovery rate (HMFDR), and Fisher statistic), maximum and mean DMR effect sizes, and annotated genes. Positive effect size indicates higher DNA methylation in males compared to females.

***Supplementary table 4-*** Overlapping genes. Genes which displayed sex-biased gene expression in GTEx and FUSION as well as sex-biased DNA methylation (according to DMRs) in the meta-analysis. Corresponding chromosome, ensemble gene ID, gene name, GTEx mash posterior effect size, GTEx local false sign rate threshold, FUSION mRNA fold change, FUSION mRNA FDR, and number of DMPs per gene. Positive effect sizes indicate higher DNA methylation or expression in males compared to females.

***Supplementary table 5-*** GO (DMPs). Gene Ontology terms identified with GSEA using the differentially methylated positions. Type of GO term-biological process (BP), molecular function (MF), and cellular component (CC), name of GO term, N represents the total number of genes in the GO term, DE represents the number of differentially methylated genes in the GO term, and SigGenesInSet are the differentially methylated genes in the GO term.

***Supplementary table 6-*** GO (DMRs). Gene Ontology terms identified with GSEA using the differentially methylated regions. Type of GO term-biological process (BP), molecular function (MF), and cellular component (CC), name of GO term, N represents the total number of genes in the GO term, DE represents the number of differentially methylated genes in the GO term, and SigGenesInSet are the differentially methylated genes in the GO term.

***Supplementary table 7-*** KEGG (DMPs). Kyoto Encyclopedia of Genes and Genomes pathways identified with GSEA using the differentially methylated positions. Description of KEGG pathway, N represents the total number of genes in the KEGG pathway, DE represents the number of differentially methylated genes in the KEGG pathway, and SigGenesInSet are the differentially methylated genes in the KEGG pathway.

***Supplementary table 8-*** KEGG (DMRs). Kyoto Encyclopedia of Genes and Genomes pathways identified with GSEA using the differentially methylated regions. Description of KEGG pathway, N represents the total number of genes in the KEGG pathway, DE represents the number of differentially methylated genes in the KEGG pathway, and SigGenesInSet are the differentially methylated genes in the KEGG pathway.

***Supplementary table 9-*** Reactome (DMPs). Reactome pathways identified with GSEA using the differentially methylated positions. Description of Reactome pathway, N represents the total number of genes in the Reactome pathway, DE represents the number of differentially methylated genes in the Reactome pathway, and SigGenesInSet are the differentially methylated genes in the Reactome pathway.

***Supplementary table 10-*** Reactome (DMRs). Reactome pathways identified with GSEA using the differentially methylated regions. Description of Reactome pathway, N represents the total number of genes in the Reactome pathway, DE represents the number of differentially methylated genes in the Reactome pathway, and SigGenesInSet are the differentially methylated genes in the Reactome pathway.

***Supplementary table 11-*** PCRs. Results from qPCR of *GGT7, FOXO3*, and *ALDH1A1* in Gene SMART cohort. Blank cells indicate sample that had a standard deviation of greater than 1 Ct between triplicates. Housekeeping genes tested are Cyclophillin, 18s rRNA, and TBP. Primer sequences and PCR conditions used.

**Supplementary table 12-.** Concentrations of circulating testosterone (nmol/L), free testosterone (pmol/L), sex hormone-binding globulin (nmol/L), and oestrogen (pmol/L) in males and females from the Gene SMART cohort. Includes 20 females (20 at rest before four weeks of exercise training, same 20 at rest after four weeks of exercise training, and six of those at rest before a control period) and 44 males (we did not have data for one male included in the DNA methylation analysis).

|  | Females, N = 46<sup>1</sup> | Males, N = 44<sup>1</sup> | p-value<sup>2</sup> |
| --- | --- | --- | --- |
| Testosterone | 1 (0) | 20 (6) | <0.001 |
| SHBG | 72 (28) | 33 (14) | <0.001 |
| Testosterone (free) | 10 (4) | 492 (160) | <0.001 |
| Estrogen | 239 (242) | 103 (24) | <0.001 |

<sup>1</sup> Mean (SD) <sup>2</sup> Welch Two Sample t-test

**Supplementary table 13-.**
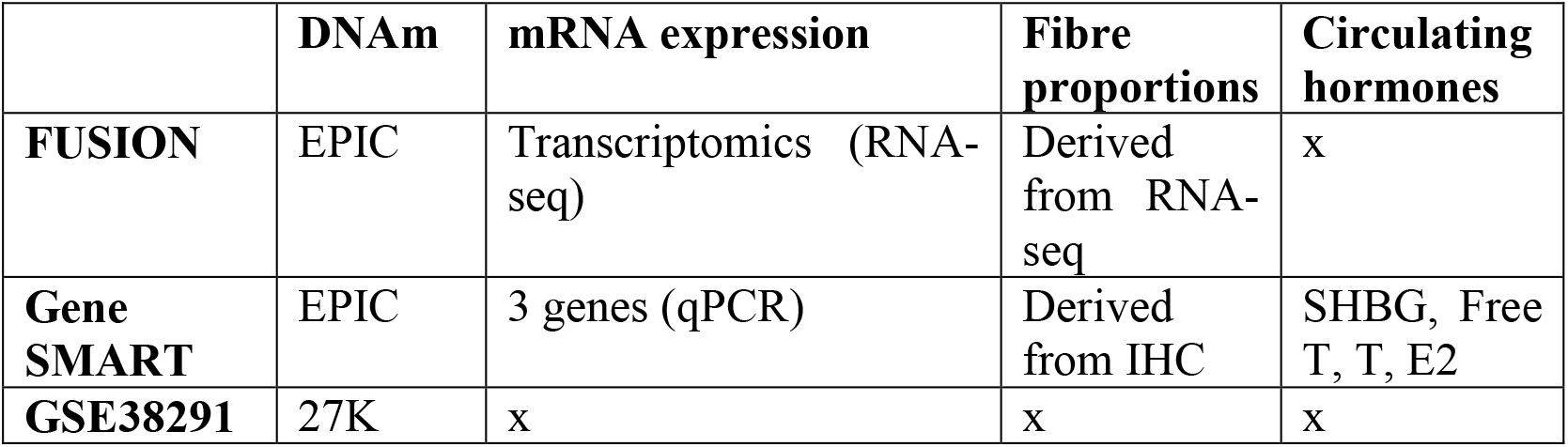
Data available for each of the datasets included in the DNA methylation meta-analysis. Immunohistochemistry (IHC); sex hormone-binding globulin (SHBG), free testosterone (Free T), testosterone (T), estradiol (E2)

***Supplementary table 14-*** List of transcription factors (TFs) included in analysis for enrichment of transcription factor binding sites (TFBSs) among differentially methylated positions (DMPs). The current UniBind database tests a total of 268 unique TFs from 518 different cell types.

## References

1. van Iterson, M., E.W. van Zwet, and B.T. Heijmans, Controlling bias and inflation in epigenome-and transcriptome-wide association studies using the empirical null distribution. Genome biology, 2017. 18(1): p. 1–13.

2. Gheorghe, M., et al., A map of direct TF DNA interactions in the human genome. Nucleic acids research, 2019. 47(4): p. e21–e21.

3. Khramtsova, E.A., L.K. Davis, and B.E. Stranger, The role of sex in the genomics of human complex traits. Nature Reviews Genetics, 2018: p. 1.

4. Takahashi, T., et al., Sex differences in immune responses that underlie COVID-19 disease outcomes. Nature, 2020. 588(7837): p. 315–320.

5. Mamlouk, G.M., et al., Sex bias and omission in neuroscience research is influenced by research model and journal, but not reported NIH funding. Frontiers in neuroendocrinology, 2020. 57: p. 100835.

6. Rawlik, K., O. Canela-Xandri, and A. Tenesa, Evidence for sex-specific genetic architectures across a spectrum of human complex traits. Genome biology, 2016. 17(1): p. 166.

7. Oliva, M., et al., The impact of sex on gene expression across human tissues. Science, 2020. 369(6509): p. eaba3066.

8. Zore, T., M. Palafox, and K. Reue, Sex differences in obesity, lipid metabolism, and inflammation—A role for the sex chromosomes? Molecular metabolism, 2018. 15: p. 35–44.

9. Arnold, A.P., Y chromosome’s roles in sex differences in disease. Proceedings of the National Academy of Sciences, 2017. 114(15): p. 3787–3789.

10. Arnold, A.P., X. Chen, and Y. Itoh, What a difference an X or Y makes: sex chromosomes, gene dose, and epigenetics in sexual differentiation, in Sex and gender differences in pharmacology. 2013, Springer. p. 67–88.

11. Golden, L.C., et al., Parent-of-origin differences in DNA methylation of X chromosome genes in T lymphocytes. Proceedings of the National Academy of Sciences, 2019. 116(52): p. 26779–26787.

12. Pihlajamaa, P., et al., Tissue-specific pioneer factors associate with androgen receptor cistromes and transcription programs. The EMBO journal, 2014. 33(4): p. 312–326.

13. Varlamov, O., C.L. Bethea, and C.T. Roberts Jr, Sex-specific differences in lipid and glucose metabolism. Frontiers in endocrinology, 2015. 5: p. 241.

14. Lopes-Ramos, C.M., et al., Sex Differences in Gene Expression and Regulatory Networks across 29 Human Tissues. Cell Reports, 2020. 31(12): p. 107795.

15. Maher, A.C., et al., Sex differences in global mRNA content of human skeletal muscle. PLoS One, 2009. 4(7): p. e6335.

16. Tarnopolsky, M., Sex differences in exercise metabolism and the role of 17-beta estradiol. Medicine & Science in Sports & Exercise, 2008. 40(4): p. 648–654.

17. Landen, S., et al., Genetic and epigenetic sex-specific adaptations to endurance exercise. Epigenetics, 2019. 14(6): p. 523–535.

18. Haizlip, K.M., B.C. Harrison, and L.A. Leinwand, Sex-Based Differences in Skeletal Muscle Kinetics and Fiber-Type Composition. Physiology (Bethesda), 2015. 30(1): p. 30–9.

19. Hunter, S.K., Sex Differences in Human Fatigability: Mechanisms and Insight to. Acta Physiol (Oxf), 2014. 210(4): p. 768–89.

20. Hall, E., et al., Sex differences in the genome-wide DNA methylation pattern and impact on gene expression, microRNA levels and insulin secretion in human pancreatic islets. Genome Biol, 2014. 15(12): p. 522.

21. Singmann, P., et al., Characterization of whole-genome autosomal differences of DNA methylation between men and women. Epigenetics & chromatin, 2015. 8(1): p. 1–13.

22. Liu, J., et al., A study of the influence of sex on genome wide methylation. PloS one, 2010. 5(4): p. e10028.

23. Davegårdh, C., et al., Sex influences DNA methylation and gene expression in human skeletal muscle myoblasts and myotubes. Stem Cell Research & Therapy, 2019. 10(1): p. 26.

24. Lindholm, M.E., et al., The human skeletal muscle transcriptome: sex differences, alternative splicing, and tissue homogeneity assessed with RNA sequencing. Faseb j, 2014. 28(10): p. 4571–81.

25. Welle, S., R. Tawil, and C.A. Thornton, Sex-Related Differences in Gene Expression in Human Skeletal Muscle. PLoS One, 2008. 3(1).

26. Gershoni, M. and S. Pietrokovski, The landscape of sex-differential transcriptome and its consequent selection in human adults. BMC Biology, 2017. 15(1): p. 7.

27. Alexander, S.E., A.C. Pollock, and S. Lamon, The effect of sex hormones on skeletal muscle adaptation in females. Eur J Sport Sci, 2021: p. 1–11.

28. Pihlajamaa, P., B. Sahu, and O.A. Jänne, Determinants of Receptor-and Tissue-Specific Actions in Androgen Signaling. Endocr Rev, 2015. 36(4): p. 357–84.

29. Varlamov, O., C.L. Bethea, and C.T. Roberts, Jr., Sex-specific differences in lipid and glucose metabolism. Front Endocrinol (Lausanne), 2014. 5: p. 241.

30. Fornes, O., et al., JASPAR 2020: update of the open-access database of transcription factor binding profiles. Nucleic Acids Res, 2020. 48(D1): p. D87–d92.

31. Lopes-Ramos, C.M., et al., Sex Differences in Gene Expression and Regulatory Networks across 29 Human Tissues. Cell Rep, 2020. 31(12): p. 107795.

32. Rajan, P., et al., Identification of novel androgen-regulated pathways and mRNA isoforms through genome-wide exon-specific profiling of the LNCaP transcriptome. PLoS One, 2011. 6(12): p. e29088.

33. Wu, Y., et al., Identification of androgen response elements in the insulin-like growth factor I upstream promoter. Endocrinology, 2007. 148(6): p. 2984–93.

34. Hevener, A.L., et al., The impact of ERα action on muscle metabolism and insulin sensitivity - Strong enough for a man, made for a woman. Mol Metab, 2018. 15: p. 20–34.

35. Wiik, A., et al., Expression of both oestrogen receptor alpha and beta in human skeletal muscle tissue. Histochem Cell Biol, 2009. 131(2): p. 181–9.

36. Vingren, J.L., et al., Effect of resistance exercise on muscle steroid receptor protein content in strength-trained men and women. Steroids, 2009. 74(13-14): p. 1033–9.

37. Mayne, B.T., et al., Large Scale Gene Expression Meta-Analysis Reveals TissueSpecific, Sex-Biased Gene Expression in Humans. Front Genet, 2016. 7: p. 183.

38. Maughan, R., J.S. Watson, and J. Weir, Strength and cross-sectional area of human skeletal muscle. The Journal of physiology, 1983. 338(1): p. 37–49.

39. Carter, S., et al., Changes in skeletal muscle in males and females following endurance training. Canadian journal of physiology and pharmacology, 2001. 79(5): p. 386–392.

40. Begue, G., et al., DNA methylation assessment from human slow-and fast-twitch skeletal muscle fibers. Journal of Applied Physiology, 2017. 122(4): p. 952–967.

41. Zierath, J.R. and J.A. Hawley, Skeletal muscle fiber type: influence on contractile and metabolic properties. PLoS biology, 2004. 2(10): p. e348.

42. Voisin, S., et al., Meta-analysis of genome-wide DNA methylation and integrative OMICs in human skeletal muscle. bioRxiv, 2020.

43. Davegårdh, C., et al., DNA methylation in the pathogenesis of type 2 diabetes in humans. Molecular metabolism, 2018. 14: p. 12–25.

44. Jones, P.A. and D. Takai, The Role of DNA Methylation in Mammalian Epigenetics. Science, 2001. 293(5532): p. 1068–1070.

45. Kundaje, A., et al., Integrative analysis of 111 reference human epigenomes. Nature, 2015. 518(7539): p. 317–330.

46. Yen, A. and M. Kellis, Systematic chromatin state comparison of epigenomes associated with diverse properties including sex and tissue type. Nature communications, 2015. 6(1): p. 1–13.

47. Voss, T.C. and G.L. Hager, Dynamic regulation of transcriptional states by chromatin and transcription factors. Nature Reviews Genetics, 2014. 15(2): p. 69–81.

48. Puig, R.R., et al., UniBind: maps of high-confidence direct TF-DNA interactions across nine species. bioRxiv, 2020.

49. Guo, S., et al., Identification of methylation haplotype blocks aids in deconvolution of heterogeneous tissue samples and tumor tissue-of-origin mapping from plasma DNA. Nature genetics, 2017. 49(4): p. 635–642.

50. VanderKraats, N.D., et al., Discovering high-resolution patterns of differential DNA methylation that correlate with gene expression changes. Nucleic acids research, 2013. 41(14): p. 6816–6827.

51. Schlosberg, C.E., N.D. VanderKraats, and J.R. Edwards, Modeling complex patterns of differential DNA methylation that associate with gene expression changes. Nucleic acids research, 2017. 45(9): p. 5100–5111.

52. Miller, A.E.J., et al., Gender differences in strength and muscle fiber characteristics. European journal of applied physiology and occupational physiology, 1993. 66(3): p. 254–262.

53. Su, J., et al., A novel atlas of gene expression in human skeletal muscle reveals molecular changes associated with aging. Skeletal muscle, 2015. 5(1): p. 1–12.

54. McCarthy, N.S., et al., Meta-analysis of human methylation data for evidence of sex-specific autosomal patterns. BMC Genomics, 2014. 15(1): p. 981.

55. Yousefi, P., et al., Sex differences in DNA methylation assessed by 450 K BeadChip in newborns. BMC genomics, 2015. 16(1): p. 911.

56. Inoshita, M., et al., Sex differences of leukocytes DNA methylation adjusted for estimated cellular proportions. Biology of sex differences, 2015. 6(1): p. 1–7.

57. Bauer, M., Cell-type-specific disturbance of DNA methylation pattern: a chance to get more benefit from and to minimize cohorts for epigenome-wide association studies. International Journal of Epidemiology, 2018. 47(3): p. 917–927.

58. Suzuki, M. and A. Bird, Suzuki MM, Bird A. DNA methylation landscapes: provocative insights from epigenomics. Nat Rev Genet 9: 465-476. Nature reviews. Genetics, 2008. 9: p. 465–76.

59. Taylor, D.L., et al., Integrative analysis of gene expression, DNA methylation, physiological traits, and genetic variation in human skeletal muscle. Proceedings of the National Academy of Sciences, 2019. 116(22): p. 10883–10888.

60. Rubenstein, A.B., et al., Single-cell transcriptional profiles in human skeletal muscle. Scientific reports, 2020. 10(1): p. 1–15.

61. De Micheli, A.J., et al., A reference single-cell transcriptomic atlas of human skeletal muscle tissue reveals bifurcated muscle stem cell populations. Skeletal muscle, 2020. 10(1): p. 1–13.

62. Domenig, S.A., A.S. Palmer, and O. Bar-Nur, Stem Cell-Based and Tissue Engineering Approaches for Skeletal Muscle Repair, in Organ Tissue Engineering. 2020, Springer International Publishing. p. 1–62.

63. Suzuki, M.M. and A. Bird, DNA methylation landscapes: provocative insights from epigenomics. Nature Reviews Genetics, 2008. 9(6): p. 465–476.

64. Pai, A.A., et al., A genome-wide study of DNA methylation patterns and gene expression levels in multiple human and chimpanzee tissues. PLoS Genet, 2011. 7(2): p. e1001316.

65. Zhou, J., et al., Tissue-specific DNA methylation is conserved across human, mouse, and rat, and driven by primary sequence conservation. BMC genomics, 2017. 18(1): p. 1–17.

66. Pöllänen, E., et al., Differential influence of peripheral and systemic sex steroids on skeletal muscle quality in pre-and postmenopausal women. Aging cell, 2011. 10(4): p. 650–660.

67. Pöllänen, E., et al., Intramuscular sex steroid hormones are associated with skeletal muscle strength and power in women with different hormonal status. Aging Cell, 2015. 14(2): p. 236–248.

68. Alexander, S.E., A.C. Pollock, and S. Lamon, The effect of sex hormones on skeletal muscle adaptation in females: Influence of sex hormones on female muscle physiology. European Journal of Sport Science, 2021(just-accepted): p. 1–27.

69. Sinha-Hikim, I., et al., Androgen receptor in human skeletal muscle and cultured muscle satellite cells: up-regulation by androgen treatment. The Journal of Clinical Endocrinology & Metabolism, 2004. 89(10): p. 5245–5255.

70. Ghanim, H., et al., Diminished androgen and estrogen receptors and aromatase levels in hypogonadal diabetic men: reversal with testosterone. Eur J Endocrinol, 2018. 178(3): p. 277–283.

71. Amar, D., et al., Time trajectories in the transcriptomic response to exercise-a meta-analysis. Nature Communications, 2021. 12(1): p. 1–12.

72. Meakin, A.S., et al., Let’s Talk about Placental Sex, Baby: Understanding Mechanisms That Drive Female-and Male-Specific Fetal Growth and Developmental Outcomes. International Journal of Molecular Sciences, 2021. 22(12): p. 6386.

73. Skinner, B.D., et al., A systematic review and meta-analysis examining whether changing ovarian sex steroid hormone levels influence cerebrovascular function. Frontiers in physiology, 2021. 12.

74. Kotamarti, V.S., et al., Risk for Venous Thromboembolism in Transgender Patients Undergoing Cross-Sex Hormone Treatment: A Systematic Review. The Journal of Sexual Medicine, 2021.

75. Qaisar, R., et al., Hormone replacement therapy improves contractile function and myonuclear organization of single muscle fibres from postmenopausal monozygotic female twin pairs. J Physiol, 2013. 591(9): p. 2333–44.

76. Lundsgaard, A.-M. and B. Kiens, Gender differences in skeletal muscle substrate metabolism-molecular mechanisms and insulin sensitivity. Frontiers in endocrinology, 2014. 5: p. 195.

77. Wijchers, P.J., et al., Sexual Dimorphism in Mammalian Autosomal Gene Regulation Is Determined Not Only by Sry but by Sex Chromosome Complement As Well. Developmental Cell, 2010. 19(3): p. 477–484.

78. De Vries, G.J., Minireview: Sex Differences in Adult and Developing Brains: Compensation, Compensation, Compensation. Endocrinology, 2004. 145(3): p. 1063–1068.

79. Ho, B., et al., X chromosome dosage and presence of SRY shape sex-specific differences in DNA methylation at an autosomal region in human cells. Biol Sex Differ, 2018. 9(1): p. 10.

80. Penny, G.D., et al., Requirement for Xist in X chromosome inactivation. Nature, 1996. 379(6561): p. 131–7.

81. Carrel, L. and H.F. Willard, X-inactivation profile reveals extensive variability in X-linked gene expression in females. Nature, 2005. 434(7031): p. 400–4.

82. Sidorenko, J., et al., The effect of X-linked dosage compensation on complex trait variation. Nature Communications, 2019. 10(1): p. 3009.

83. Tukiainen, T., et al., Landscape of X chromosome inactivation across human tissues. Nature, 2017. 550(7675): p. 244–248.

84. Grafodatskaya, D., et al., Multilocus loss of DNA methylation in individuals with mutations in the histone H3 lysine 4 demethylase KDM5C. BMC Med Genomics, 2013. 6: p. 1.

85. Trolle, C., et al., Widespread DNA hypomethylation and differential gene expression in Turner syndrome. Scientific reports, 2016. 6(1): p. 1–14.

86. Sharma, A., et al., DNA methylation signature in peripheral blood reveals distinct characteristics of human X chromosome numerical aberrations. Clinical epigenetics, 2015. 7(1): p. 1–15.

87. Villicaña, S. and J.T. Bell, Genetic impacts on DNA methylation: research findings and future perspectives. Genome biology, 2021. 22(1): p. 1–35.

88. Smith, J., et al., Promoter DNA hypermethylation and paradoxical gene activation. Trends in cancer, 2020. 6(5): p. 392–406.

89. Mielcarek, M., et al., HDAC4 as a potential therapeutic target in neurodegenerative diseases: a summary of recent achievements. Frontiers in cellular neuroscience, 2015. 9: p. 42.

90. Pigna, E., et al., HDAC4 preserves skeletal muscle structure following long-term denervation by mediating distinct cellular responses. Skeletal muscle, 2018. 8(1): p. 1–16.

91. McGee, S.L., et al., Exercise-induced histone modifications in human skeletal muscle. The Journal of Physiology, 2009. 587(24): p. 5951–5958.

92. Meng, Z.-X., et al., Glucose sensing by skeletal myocytes couples nutrient signaling to systemic homeostasis. Molecular cell, 2017. 66(3): p. 332–344. e4.

93. Brown, W.M., Exercise-associated DNA methylation change in skeletal muscle and the importance of imprinted genes: a bioinformatics meta-analysis. Br J Sports Med, 2015. 49(24): p. 1567–78.

94. Stefanetti, R.J., et al., Recent advances in understanding the role of FOXO3. F1000Research, 2018. 7.

95. Bui, T.T., et al., γ-Glutamyl transferase 7 is a novel regulator of glioblastoma growth. BMC Cancer, 2015. 15: p. 225.

96. Skelly, L.E., et al., Effect of sex on the acute skeletal muscle response to sprint interval exercise. Experimental physiology, 2017. 102(3): p. 354–365.

97. Welle, S., et al., Skeletal muscle gene expression profiles in 20–29 year old and 65–71 year old women. Experimental gerontology, 2004. 39(3): p. 369–377.

98. Baldelli, S., et al., Glutathione and Nitric Oxide: Key Team Players in Use and Disuse of Skeletal Muscle. Nutrients, 2019. 11(10).

99. Yan, X., et al., The gene SMART study: method, study design, and preliminary findings. BMC Genomics, 2017. 18(Suppl 8): p. 821.

100. Ribel-Madsen, R., et al., Genome-wide analysis of DNA methylation differences in muscle and fat from monozygotic twins discordant for type 2 diabetes. PloS one, 2012. 7(12): p. e51302.

101. Tian, Y., et al., ChAMP: updated methylation analysis pipeline for Illumina BeadChips. Bioinformatics, 2017. 33(24): p. 3982–3984.

102. Pidsley, R., et al., Critical evaluation of the Illumina MethylationEPIC BeadChip microarray for whole-genome DNA methylation profiling. Genome biology, 2016. 17(1): p. 1–17.

103. Chen, Y.-a., et al., Cross-reactive DNA microarray probes lead to false discovery of autosomal sex-associated DNA methylation. The American Journal of Human Genetics, 2012. 91(4): p. 762–764.

104. Leek, J.T., et al., Package ‘sva’. 2014.

105. Leek, J.T., et al., Tackling the widespread and critical impact of batch effects in high-throughput data. Nature Reviews Genetics, 2010. 11(10): p. 733–739.

106. Willer, C.J., Y. Li, and G.R. Abecasis, METAL: fast and efficient meta-analysis of genomewide association scans. Bioinformatics, 2010. 26(17): p. 2190–2191.

107. Smyth, G.K., limma: Linear Models for Microarray Data, in Bioinformatics and Computational Biology Solutions Using R and Bioconductor, R. Gentleman, et al., Editors. 2005, Springer New York: New York, NY. p. 397–420.

108. Benjamini, Y. and Y. Hochberg, Controlling the false discovery rate: a practical and powerful approach to multiple testing. Journal of the royal statistical society. Series B (Methodological), 1995: p. 289–300.

109. Benjamin, D.J., et al., Redefine statistical significance. Nature Human Behaviour, 2018. 2(1): p. 6.

110. Peters, T.J., et al., De novo identification of differentially methylated regions in the human genome. Epigenetics & chromatin, 2015. 8(1): p. 6.

111. Ferrari, S. and F. Cribari-Neto, Beta regression for modelling rates and proportions. Journal of applied statistics, 2004. 31(7): p. 799–815.

112. Van Buuren, S. and K. Groothuis-Oudshoorn, Multivariate imputation by chained equations in RJ Stat. Softw.

113. Zhou, W., P.W. Laird, and H. Shen, Comprehensive characterization, annotation and innovative use of Infinium DNA methylation BeadChip probes. Nucleic acids research, 2017. 45(4): p. e22–e22.

114. Fishilevich, S., et al., GeneHancer: genome-wide integration of enhancers and target genes in GeneCards. Database, 2017. 2017.

115. Phipson, B., J. Maksimovic, and A. Oshlack, missMethyl: an R package for analyzing data from Illumina’s Human Methylation45O platform. Bioinformatics, 2016. 32(2): p. 286–288.

116. Maksimovic, J., A. Oshlack, and B. Phipson, Gene set enrichment analysis for genome-wide DNA methylation data. bioRxiv, 2020.

117. Schiaffino, S. and C. Reggiani, Fiber types in mammalian skeletal muscles. Physiol Rev, 2011. 91(4): p. 1447–531.

118. Bloemberg, D. and J. Quadrilatero, Rapid determination of myosin heavy chain expression in rat, mouse, and human skeletal muscle using multicolor immunofluorescence analysis. PloS one, 2012. 7(4): p. e35273.

119. Godsland, I.F., et al., The Effects of Different Formulations of Oral Contraceptive Agents on Lipid and Carbohydrate Metabolism. New England Journal of Medicine, 1990. 323(20): p. 1375–1381.

120. Urbut, S.M., et al., Flexible statistical methods for estimating and testing effects in genomic studies with multiple conditions. Nature genetics, 2019. 51(1): p. 187–195.

